# Evaluation of hemispherical photos extracted from smartphone spherical panorama images to estimate canopy structure and forest light environment

**DOI:** 10.1101/2020.12.15.422956

**Authors:** A. Z. Andis Arietta

## Abstract

Hemispherical photography (HP) is one of the most commonly employed methods to estimate forest canopy structure and understory light environments. Traditional methods require expensive, specialized equipment, are tedious to deploy, and are sensitive to exposure settings. In contrast, modern smartphone cameras are readily available and make use of ever-improving software to produce images with high dynamic range and clarity, but lack suitable hemispherical lenses. Thus, despite the fact that almost all ecologists and foresters carry a high-powered, image processing device in our pockets, we have yet to fully employ it for the purpose of data collection. As an alternative, hemispherical images can be extracted from spherical panoramas produced by many smartphone camera applications. I compared hemispherical photos captured with a digital single lens reflex camera and 180° lens to those extracted from smartphone spherical panoramas (SSP) for 72 sites representing a range of canopy types and densities. I estimated common canopy and light measures (canopy openness, leaf area index, and global site factor) as well as image quality measures (total gap area, number of gaps, and relative gap size) to compare methods. The SSP HP method leverages built-in features of current generation smartphones including exposure metering over restricted field-of-view, high dynamic range tonal correction, computational sharpening, high pixel density, and automatic leveling via the phone’s built-in gyroscope to yield an accurate alternative to traditional HP in canopy estimation. Although the process of stitching together multiple photos occasionally produces artifacts in the SSP HP images, estimates of canopy openness and global site factor are highly correlated with those of traditional methods (R^2^ > 0.9) and are comparable to under- or over-exposing traditional HP by 1-1.5 stops. In addition to superior image quality, SSP HP requires no additional equipment or exposure settings and is likely to prove more robust to uneven lighting conditions by avoiding wide-angles lenses and exploiting HDR images.

## 1. Introduction

Ecological patterns and processes in forests are mediated by the canopy in critical ways. The structure of the canopy directly alters the below-canopy light regime and microclimate which indirectly impacts microhabitat features such as soil moisture, snowpack, and understory plant community (Jennings, Brown and Sheil, 1999). Because of its fundamental importance, forest managers and ecologists alike require methods to accurately quantify canopy structure and light environments.

In recent decades, digital hemispherical photography (HP) has arisen as the most popular sampling method for canopy measurement, primed by advances in digital photography equipment and software for image analysis (Promis, 2013; Chianucci, 2020). This technique uses extreme wide-angle lenses with a field-of-view (FOV) of 180° that projects an entire hemisphere of view onto the camera sensor, resulting in a circular hemispherical image (Rich, 1990). Pixels within the image are then classified into binary sky (white) or canopy elements (black) manually or algorithmically using global or local thresholds (Glatthorn and Beckschäfer, 2014). From the binarized images, canopy structure measures such as canopy openness, leaf area index, gap fraction, etc. can be estimated (Frazer, Trofymow and Lertzman, 1997; Gonsamo, D’odorico and Pellikka, 2013; Chianucci, 2020). By plotting a sun path onto the binarized image the canopy structure can be used to infer understory light regimes, or Site Factors, given estimates of prevailing above-canopy direct and diffuse radiation (Anderson, 1964). Light values can be integrated over time to yield seasonal estimates of light environments from a single sampling event (Frazer, Trofymow and Lertzman, 1997)(or at least two samples in deciduous canopies (e.g. Halverson *et al.*, 2003)). Thus, HP offers an efficient, non-destructive method of estimating forest microhabitat features.

However, HP suffers from methodological drawbacks relating to the difficulty of capturing images in field settings and the sensitivity of estimates to variation in image acquisition (Beckschäfer *et al.*, 2013; Bianchi *et al.*, 2017). Traditionally, HP requires a single lens reflex camera (or more commonly nowadays, digital single lens reflex (DSLR)) equipped with a specialized hemispherical lens and self-leveling tripod. For the most accurate estimates, images must be acquired against uniformly overcast skies or the fleeting light at dusk or dawn with the camera level to the horizon and with closely calibrated exposure. The reliance on DSLRs stems from the need to manually fine-tune exposure and the advantage of large sensors. Large sensors record more pixels per area of view and more information per pixel, which translates to more accurate classification of canopy gaps. Careful tuning of exposure settings is critical to avoid major inaccuracies in final estimates (Zhang, Chen and Miller, 2005; Beckschäfer *et al.*, 2013).

Typical camera and hemispherical lens systems are expensive and challenging to deploy in the field. In response, researchers have attempted to develop new methods including using smartphones with clip-on lenses (Tichý, 2016; Bianchi *et al.*, 2017) and eschewing specialized leveling devices (Origo *et al.*, 2017), to varying success. Smartphones are limited by small sensors and the low quality of aftermarket fisheye lenses. Other researchers have developed methods that use standard cameras with reduced FOV lenses and account for non-hemispherical images (i.e. restricted view photography) to estimate canopy structure with less sensitivity to camera exposure while maximizing the full frame of the sensor (Chianucci, 2020). Yet, without a full hemisphere of view, this method cannot be used for estimating light environments. Thus, despite the fact that almost all ecologists and foresters carry a high-powered, image processing device in our pockets, we have yet to fully employ it for the purpose of data collection.

Here, I test a novel method of HP acquisition from smartphone spherical panoramas (SSP HP) for estimating canopy measures. This method leverages the advantages of restricted view photography and the utility of smartphones to produce true circular HP at higher resolution than traditional DSLR HP without the need for additional equipment (i.e. leveling device or lens). I compare estimates from HP images extracted from SSP HP to traditional DSLR HP images and an alternative smartphone method with a fisheye lens proposed by Bianchi et al. (2017).

Spherical panoramas can be generated on any modern smartphone with pre-installed software like Google Camera (Google LLC), or free applications like Google Street View (Google LLC) (available for Android OS or iOS). With Google Camera, spherical panoramas are composed from 36 individual images with restricted FOV (57° FOV with Google Pixel 4a). Images are taken by rotating the camera around a central point, guided by the camera’s spatial mapping and aided by the device’s internal gyroscope and compass. The smartphone camera software automatically merges the individual images into a spherical projection using interest point detection and scale invariant feature transformation to accommodate imperfect viewing distance and viewing angle between images (Szeliski and Shum, 1997; Brown and Lowe, 2007). Spherical panoramas are recorded in equirectangular projection from which the top half can be easily remapped into polar projection as a circular HP (Fig. 1).

**Figure 1.**
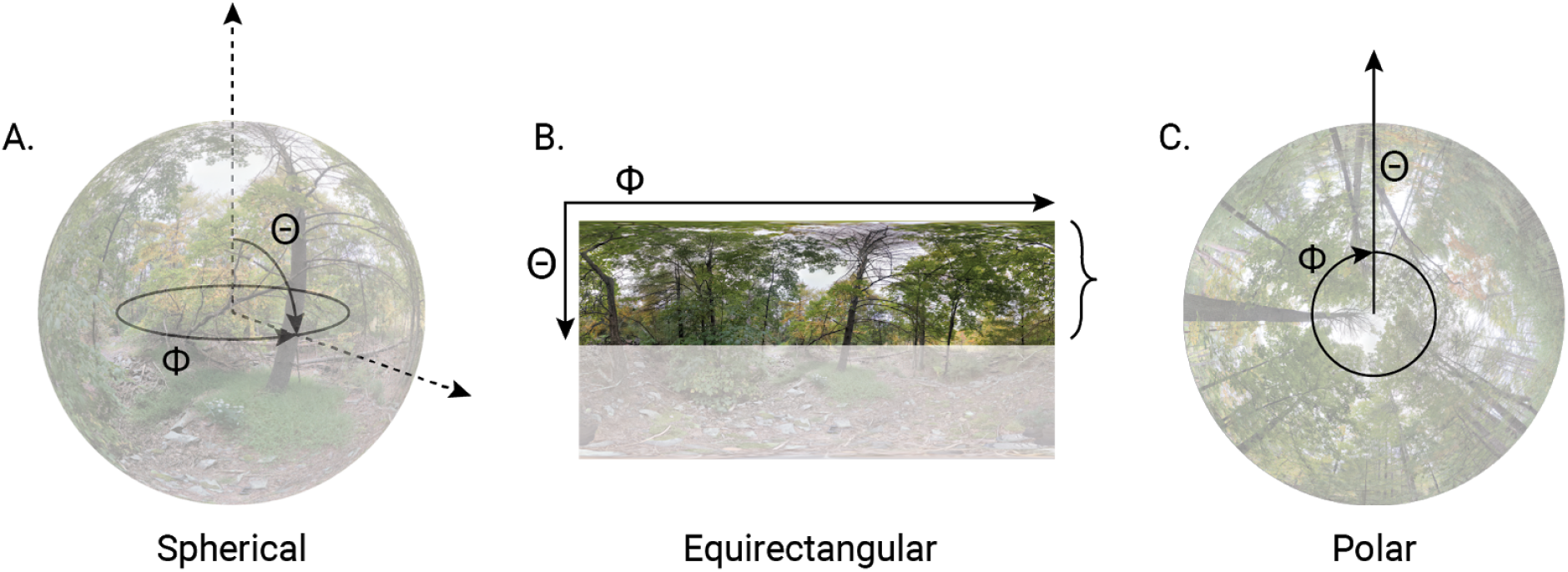
Spherical panoramas (A) are stored and output from smartphones as 2D images with equirectangular projection (B). Because spherical panoramas are automatically leveled using the phone gyroscope, the top half of the equirectangular image corresponds to the upper hemisphere of the spherical panorama. The top portion of the equirectangular image (B) can then be remapped onto the polar coordinate plane to create a circular hemispherical photo (C). In all images, zenith and azimuth are indicated by Θ and Φ, respectively.

Modern smartphones overcome physical limitations of small optics and sensors by employing computational photography techniques that merge multiple images to create a single, high resolution image (Barbero-García *et al.*, 2018). The result is a composite image that retains the sharpest elements and most even exposure of each individual photo that is sharper (Gunturk, 2017) and with greater dynamic range than any individual photo (Lukac, 2017). Modern computational photography with small sensors can rival images produced by much larger, DSLR-sized sensors (Ignatov *et al.*, 2017).

## 2. Methods

I estimated canopy structure and light values from two sources: hemispherical photos captured with a DSLR camera (Canon 60D; Canon Inc., Tokyo, Japan) equipped with a circular hemispherical lens (Sigma 8mm f3.5 EX DG; Sigma Corp., Ronkonkoma, N.Y., USA) and spherical panoramic images captured with a smartphone (Pixel 4a; Google LLC, Menlo Park, C.A., USA) and native spherical panorama software (Google Camera v.8.1.011.342784911). In addition, I simulated images to approximate the method proposed by Bianchi et al. (2017).

I acquired images from 35 sites at Yale Preserve, New Haven, Connecticut, USA and 37 sites at Rockstock property, Woodstock, New York, USA on 4 July 2020 and 27 September 2020, respectively. Photo sites were selected to represent a distribution of canopy species (deciduous, coniferous, and mixed), openness, and gap size. All images were captured at breast height (approx. 1.3 m) on uniformly overcast days with smartphone images taken immediately following each DSLR image.

### 2.1 DSLR protocol

Prior to image acquisition, I established exposure settings two stops overexposed relative to open sky following Brown et al. (2000) and Beckchafer et al. (2013) with maximum ISO values of 1000 and minimum shutter speed of 1/100 s (Chianucci and Cutini, 2012). Images were recorded in Canon RAW format (.CR2) with the camera oriented perpendicular to gravity enabled by a dual-axis gimbal and with the top of the image oriented toward magnetic north.

Even on uniformly overcast days, the sky brightness changes over time. To account for this, I adjusted white point values in Adobe Lightroom 5.7.1 to ensure that the grey value of the brightest sky pixels aligned to full white (Beckschäfer *et al.*, 2013) and exported the images as full-resolution (5184 x 3456 pixels) JPEG files. These images are considered the standard reference for comparison throughout further analysis.

To compare the discrepancy due to image acquisition methods to the discrepancy due to incorrect exposure, I additionally created output files with exposure values adjusted 1, 2, 3, 4, or 5 values above and below the original exposure in Adobe Lightroom. I included a circular mask along the perimeter of the circular image to prevent glare at the margins from impacting downstream estimates.

### 2.2 Smartphone spherical panorama protocol

I created spherical panoramas using a Pixel 4a smartphone—Google’s mid-range consumer-grade smartphone model—with the Google Camera application. Spherical panoramas are composed from 36 overlapping images of the entire 360 degree field of view. I began each spherical image sequence facing toward magnetic north (azimuth 0°; this becomes the top of the circular hemispherical image after processing). For this study, I ascertained a northern heading with an external compass first for DSLR HP images and used this to orient the SSP HP. However, the yaw angle from the metadata of the resulting SSP HP image can be used to rotate the image in post-processing to the proper orientation regardless of the direction of capture (Li and Ratti, 2019). The first image of the panorama must be taken with the phone levelled to the horizon; the camera software facilitates this by placing a dot on the screen and disallowing images with too great of pitch or roll to the camera. Subsequent images are similarly guided by on-screen targets. Twelve images centered along the horizon comprise zenith 72° to 108°. Nine images in the upper and lower hemisphere cover zenith 36° to 72° or 108° to 144°, respectively. Similarly, three images each comprise the remaining area at the poles. Although no order is specified by the application, I followed the same capture order for every spherical panorama, first rotating to capture the twelve images sequentially around the horizon. Next, I sequentially captured the nine images for zenith 36° to 72°, followed by three images for zenith 0° to 36°. I then followed the same order in the lower sphere. The phone’s internal gyroscope is used to automatically level the horizon of the sphere. Care must be taken to rotate and pan the smartphone camera treating the camera as the centerpoint, rather than rotating the camera around one’s body.

Spherical panoramas are created by tiling multiple planar images into a geodesic polyhedron and then stitching the images into a spherical, omnidirectional image which can be viewed in 3D (Fangi and Nardinocchi, 2013). The spherical images are mapped into two dimensions following an equirectangular projection in which the zenith angle corresponds to the rectangular y-axis and azimuth angle corresponds to the rectangular x-axis (Fangi and Nardinocchi, 2013) (Fig. 1). Conveniently, when the sphere is levelled to the horizon, the top half of the rectangular panoramic image depicts the upper hemisphere of view and can be cropped and remapped into a polar projection (e.g. Li and Ratti, 2019).

I extracted the top half of the equirectangular panorama JPEG file (i.e. the top hemisphere) and converted it to a circular hemispherical image via polar projection in GIMP (Gnu Image Manipulation Program v.2.10.20) with batch processing implemented in BIMP (Batch Image Manipulation Plugin v.2.4) (setting files are included in the data archive along with scripts for an alternative processing from the command line with ImageMagick v.7.0.10).

Transformation from equirectangular to polar projection with square pixels requires either downsampling pixels closer to the pole or interpolating pixels near the horizon, or both. I retained the width of the equirectangular image in the transformation to polar projection, resulting in a SSP HP image with diameter equal to the width of the equirectangular projection. Thus, the circumference of the HP at zenith 57° degrees (~ 1 radian), is equal to the width of the equirectangular image. Pixels circumscribing zenith angles greater or less than 57° are upscaled or downsampled, respectively. The area from zenith 0° to 57° is important for canopy estimates as gap fraction measurements in this portion of the hemisphere are insensitive to leaf inclination angle, allowing for estimation of LAI without accounting for leaf orientation. The result is a true HP with over 900% of the resolution of traditional DSLR HPs. Because the area of the SSP HP is larger than the upper half of the equirectangular image (i.e. more upscaling than downscaling), I test the impact of resolution below.

Unlike DSLR images, the white point does not need to be adjusted for smartphone images as this is automatically controlled by the device’s high dynamic range (HDR) routine. However, the HDR routine yields a more homogenous histogram and consolidates pixel values at the mid-tones (Fig. 2), which can make it difficult for binarization algorithms to differentiate between sky and canopy pixels. Contrast-stretching can facilitate pixel classification (Macfarlane *et al.*, 2014). To test the effect of contrast-stretching, I output two sets of SSP HP with and without expanding the tonal range by 5 (2%) in GIMP prior to polar projection conversion.

**Figure 2.**
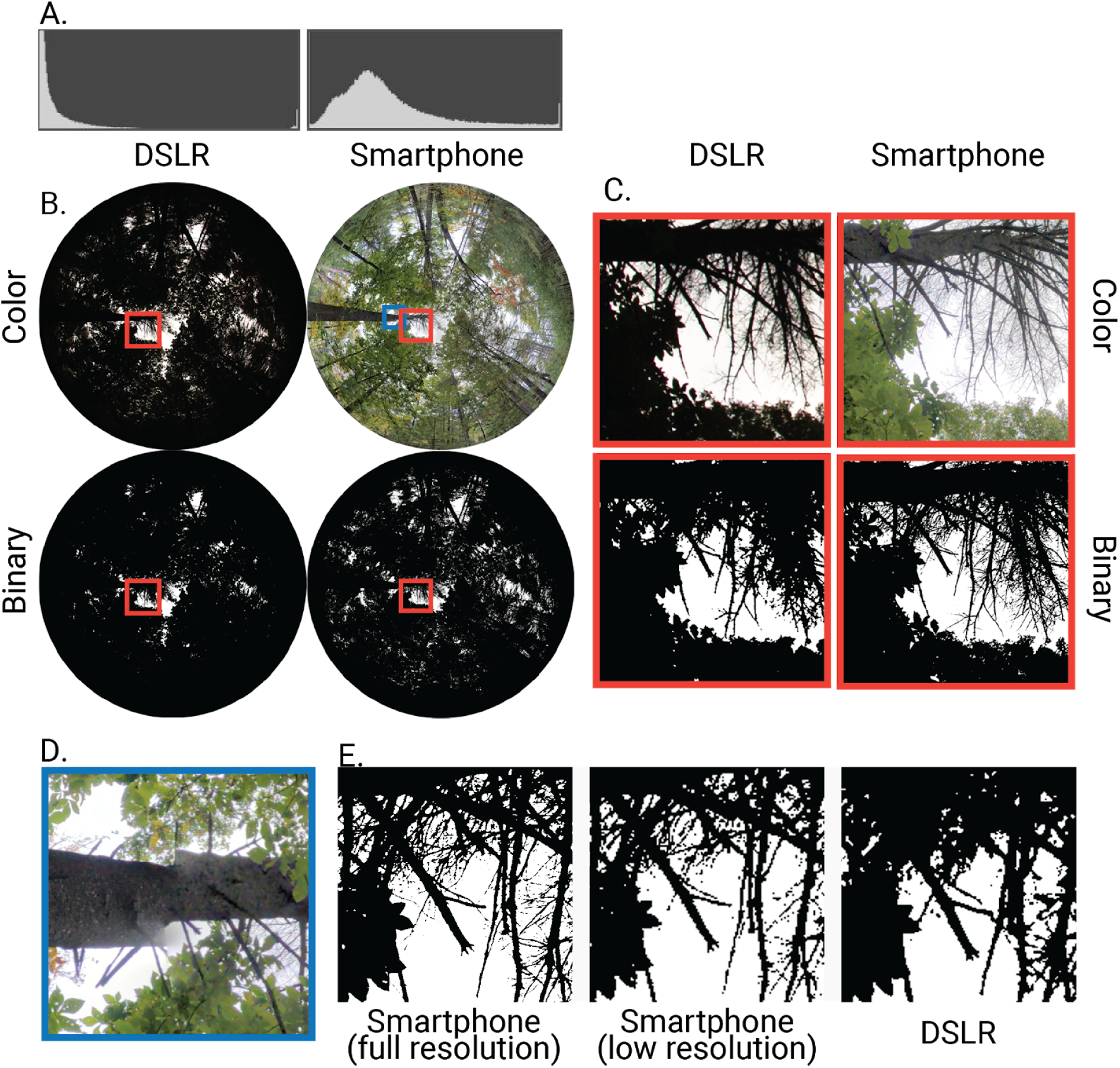
Comparisons of smartphone spherical panorama hemispherical photographs (SSP HP) (right B and C) to traditional DSLR hemispherical photographs (DSLR HP) (left B and C) captured at the same site. Details of the same subsection of the canopy, indicated by orange boxes, are expanded in C. Binarized images are shown below color images in B and C. Image histograms differ in the distribution of luminance values in the blue color plane (A). In panel E, a section of the canopy from full resolution SSP HP (left), downsampled SSP HP (middle), and DSLR HP (right) is further expanded to demonstrate the effect of image clarity on pixel classification. An example of an incongruous artifact resulting from misalignment in the spherical panorama is outlined in blue in A and expanded in D.

Circular hemispherical images from DSLR photos (diameter: 2885 p; area: 6.5 MP) are considerably smaller than those produced from photo spheres (diameter: 8704 p; area: 59.5 MP). Resolution can impact canopy estimate because larger portions of the hemisphere are averaged into each pixel, leading to higher proportion of mixed pixels and underestimates of small gaps (Macfarlane, 2011). To test the impacts of the resolution gain of the SSP HP, I exported an additional set of images downsampled to match the diameter of the DSLR photos (2885 p) in GIMP.

### 2.3 Fisheye lens simulation

Bianchi et al. (2017) proposed a method of approximating hemispherical photos from two perpendicular smartphone images using a fisheye lens adapter with 150° diagonal FOV. In order to compare this technique to the SSP HP method proposed in this study, I used the SSP HP to simulate images captured from two perpendicular Pixel 4a photos. I downsampled the SSP HP to 6049 p diameter and applied a black mask that simulates Pixel 4a image dimensions (5802 x 4352 p) with 150° FOV.

## 3. Analysis

### 3.1 Binarization and canopy estimates

The processing steps above yielded 16 sets of HP images in JPEG format (Fig. S1): standard DSLR HP (no exposure adjustment), ten sets of exposure-adjusted DSLR HP, four sets of SSP HP at full or low resolution with or without contrast adjustment, and one set of fisheye HP with appropriate contrast adjustments. From this point, all images received the same processing steps.

I binarized images with the Hemispherical 2.0 plugin (Beckschäfer, 2015) for ImageJ v.1.51k which uses the “Minimum” algorithm (Prewitt and Mendelsohn, 1966) applied to the blue color channel of the image to automatically classify pixels and output binary images in TIFF format (Beckschäfer, 2015). During binarization, the program estimates total gap area and number of gaps, which I recorded for further analysis. I converted the binary TIFF files to BMP format in batch with ImageJ.

I used Gap Light Analyzer v.2.0 (Frazer, Canham and Lertzman, 1999) to estimate additional canopy structure measures and light transmittance. I used identical configuration parameters for all image sets with two exceptions. One, I adjusted the lens projection parameters per camera device. Two, I adjusted coordinates, elevation, and declination per location. HP images created from spherical panoramas conform to true polar projection whereas the Sigma hemispherical lens used for DSLR HP images conforms to an equisolid projection (see parameter and lens configuration files in data archive). Gap Light Analyzer is implemented in a graphical interface without an option for command line input. So, I wrote a custom macro-script in AutoHotKey v.1.1.33.02 to batch process the images (script available in the data archive). I recorded canopy openness, leaf area index, and global site factor measures from Gap Light Analyzer for further analysis.

### 3.2 Statistical analysis

All statistical analyses were performed in R v.3.6.2 (R Core Team, 2019). I used ordinary least squares regression to compare differences between multiple image sets. I focused on three HP image characteristics--number of gaps, total gap area, and relative gap size--to evaluate the difference in image quality between methods. I focus on three canopy measures--canopy openness (CO), effective leaf area index (LAI), and global site factor (GSF)--to evaluate the similarity of canopy estimates between methods. Gap fraction, the ratio of white to black pixels, is the most foundational measure of HP. Canopy openness is similar to gap fraction, but weights pixels by zenith angle and is a more appropriate measure when comparing HP with different lens distortion which bias gap size at different zenith angles (Frazer, Trofymow and Lertzman, 1997; Gonsamo, D’odorico and Pellikka, 2013). I calculated relative gap size as the area of the mean canopy gap, standardized to the total image size and is equal to CO divided by the number of gaps. LAI is a comparison of leaf area relative to horizontal ground area integrated over zenith angles 0 to 60 degrees (Chianucci, 2020). GSF is a measure of canopy radiation as a weighted average of the proportion of direct and indirect radiation transmitted through the canopy to that above the canopy (Anderson, 1964). The three canopy measures are a function of the image quality measures and represent the range of the inference for which researchers use HP.

I made a series of inferences defining various sets of images as reference or comparison. First, I used the standard DSLR HP images to determine if canopy measures differed between sites from the two locations by treating location as an independent variable and each canopy measure as dependent variables, respectively. Second, I considered the effect of contrast adjustments to reduce erroneous binarization outliers by comparing full resolution and downsampled SSP HP with and without contrast-stretching (comparison sets) to the standard DSLR HP set (reference set). Third, I compared the effects of downsampling on SSP HP images by treating the full resolution images as reference and calculating the percent difference in canopy measures from the low resolution SSP HP set.

Additionally, I tested for differences between all image sets by treating the standard DSLR HP images as reference. I computed percent differences and fit linear regression models with the comparison image set as independent variables. Thus, the effect size of the dissimilarity between image sets is measured by the magnitude of the deviation of the slope from 1 or the intercept from 0 and the correlation coefficient indicates the appropriateness of the comparison as an alternative approximation of the reference. In addition, I compare canopy estimates between SSP HP and standard DSLR HP, using variability among exposure settings of DSLR HP as a qualitative measure of the effect size of the difference in methods. Finally, I compared SSP HP generated from spherical panoramas to HP generated from smartphone images captured with a fisheye lens.

## 4. Results

The canopies surveyed in this study ranged from densely closed (CO_min_ = 1%, GSF_min_ = 1) to moderately open (CO_max_ = 40%, GSF_max_ = 63). Most sites skew toward denser canopies (CO_IQR_ = 2% - 6%, GSF_IQR_ = 3 - 9). This is advantageous for the purpose of this study as estimates from dense canopies are substantially more sensitive to HP settings (Beckschäfer *et al.*, 2013). There was no significant difference between locations (all p > 0.08; Table S1).

HP images generated from spherical panoramas were noticeably sharper than those captured with a DSLR and hemispherical lens (Fig. 2C & 2E). The greater sharpness resulted in more definition of fine canopy structure even when scaled to the same resolution (Fig 2E). Close inspection of the smartphone HP reveals occasional infidelities arising from the process of stitching the panoramas (Fig. 2D). These artifacts appear as discontinuities or overlapping of canopy elements.

### 4.1 Effect of contrast-stretching on SSP HP

Unlike most DSLR HP images, which exhibit bimodal distribution of pixel tones, the HDR pre-processing of smartphone cameras produces images with tonal values with a normal distribution centered around the midpoint (Fig. 2A). This poses a problem for binarization algorithms that iteratively seek a global minimum along the histogram and can lead to extreme over-classification of sky pixels at lower resolutions or restricted FOV. One full resolution SSP HP image (1%) and three low resolution SSP HP images (4%) were incorrectly classified. Twelve fisheye HP images (17%) were incorrectly classified. I applied a 2% contrast-stretch to all SSP HP images and up to 8% as needed for fisheye HP images to ensure no misclassifications. Only the contrast adjusted images were retained for further analysis.

### 4.2 Effect of resolution on SSP HP

Downsampling SSP HP images from 59.5 MP to 6.5 MP (−815%) to match the resolution of DSLR HP images resulted in nearly half the number of gaps (−48%) of larger relative size (+91%) compared to the full resolution SSP HP images (Table 1). Downsampling had minimal effect on canopy structure and light estimates, however (Table 1). Downsampling resulted in only a 1% decrease in CO and GSF compared to full resolution SSP HP images. Downsampling had a minimal effect of increased LAI (1%). Reduction in resolution minorly increased variance in CO (1%), GSF (1%), and LAI (4%), but decreased the variance in number of gaps (−61%) and gap area (−89%) considerably. Downsampled images exhibited greater variance in relative gap size (+107%).

**Table 1.** Comparison of SSP HP images at full resolution and low resolution when downsampled to the same area as DSLR HP images.

|  |  |  |  |  |
| --- | --- | --- | --- | --- |
| Measure | Full Resolution<br>Mean (S.D.) | Low Resolution<br>Mean (S.D.) | Mean % Dif. | S.D. % Dif. |
| Canopy openness (%) | 6.78 (5.35) | 6.7 (5.41) | -1.15 | 0.96 |
| GSF | 9.69 (8.95) | 9.56 (8.99) | -1.31 | 0.48 |
| LAI | 3.19 (0.59) | 3.23 (0.62) | 1.01 | 4.25 |
| Gap area (MP) | 4.02 (3.18) | 0.44 (0.36) | -89.16 | -88.95 |
| No. of gaps (x100) | 245.32 (50.75) | 128.16 (19.82) | -47.76 | -60.96 |
| Gap size (relative) | 0 (0) | 0 (0) | 91.01 | 106.89 |

### 4.3 Comparison to DSLR HP

Full resolution SSP HP retained more gaps (591%) and smaller relative gap size (−74%) compared to DSLR HP (Table S2). Although downsampling reduces the difference, SSP HP images with the same resolution as DSLR HP exhibit more (261%) gaps of smaller relative size (−53%) (Table S2). The difference in relative gap size increases with greater canopy openness (Fig. 3). Despite the larger difference in gap number, the total gap area in downsampled smartphone images was just 51% larger than DSLR HP reference images (Table S2).

**Figure 3.**
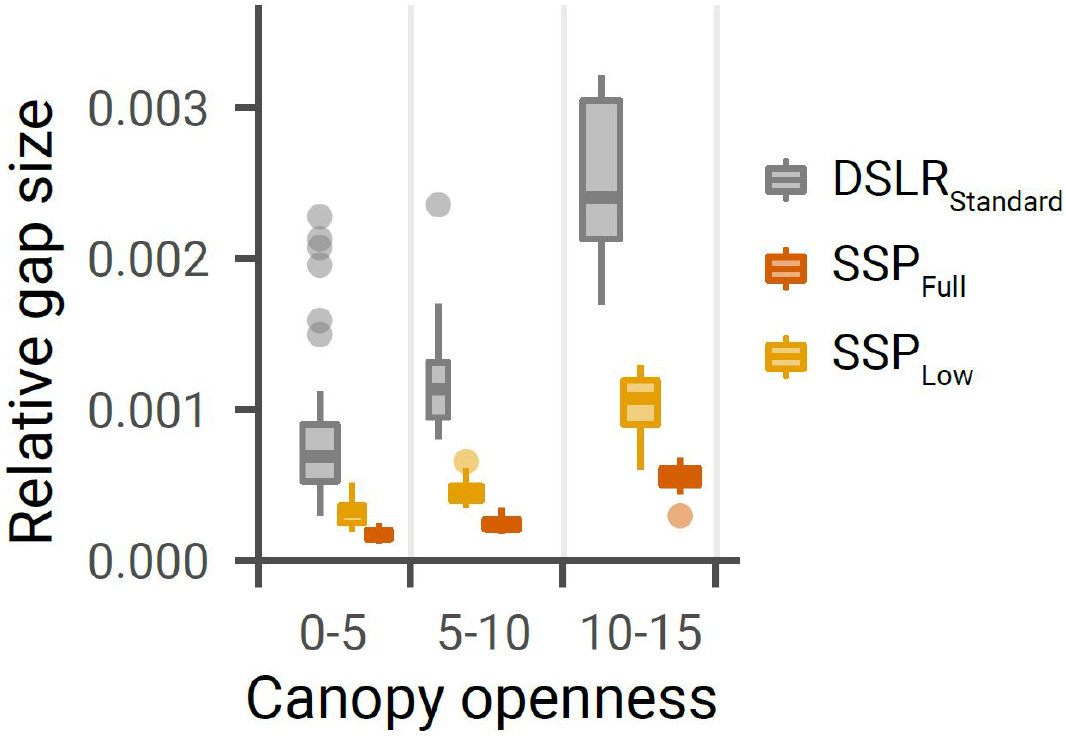
Relative gap size related to canopy openness for standard DSLR HP (grey), downsampled SSP HP (light orange), and full resolution SSP HP (dark orange). Values were binned into three canopy openness thresholds (0-5, 5-10, 10-15). Values greater than CO 15 were excluded due to lack of sites with high values.

Estimates from SSP HP were greater for CO (Full = +63%, Low = +60%), greater for GSF (Full = +57%, Low = 53%), and lower for LAI (Full = −21%, Low = −20%) compared to the reference DSLR HP (Fig. 4, Table S3). These differences are comparable to the effects of over-exposing DSLR images by 1 to 1.5 stops. SSP HP estimates of CO and GSF were highly correlated with DSLR HP estimates (R^2^ > 0.9) while LAI was moderately correlated (R^2^ = 0.64) (Fig. 5, Table S4).

**Figure 4.**
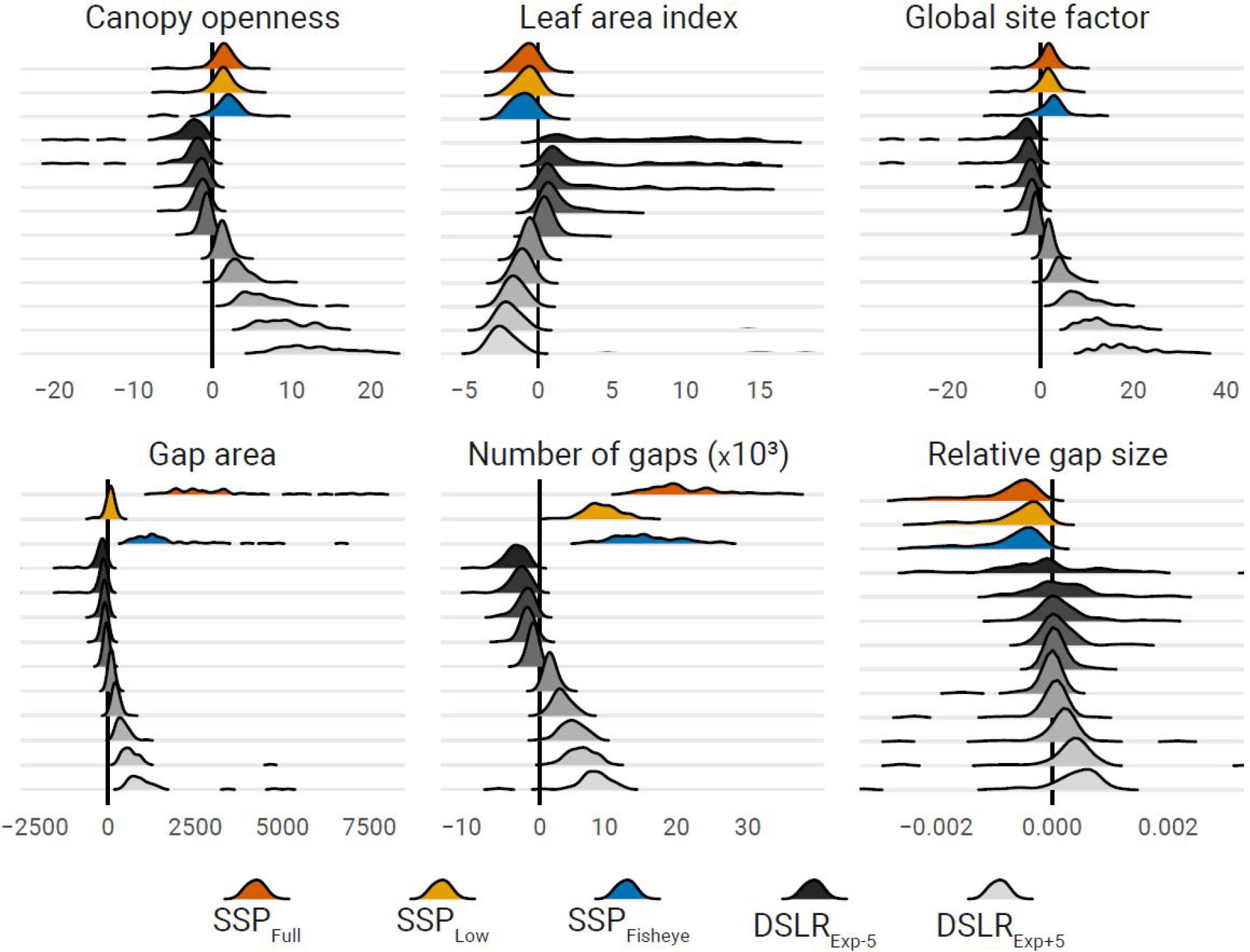
Difference in canopy structure and light environment estimates between reference (standard DSLR HP) and full resolution SSP HP (dark orange), low resolution SSP HP downsample to match the standard DSLR resolution (light orange), fisheye HP (blue), and DSLR HP with exposure adjusted from +5 to −5 (light to dark). SSP HP images were generated from spherical panoramas taken with Google Pixel 4a and Google Camera. Fisheye HP images were simulated from smartphone HP for two intersecting 150° FOV images from a Pixel 4a. DSLR HP were captured with Canon 60D and Sigma 4.5mm f2.8 hemispherical lens.

**Figure 5.**
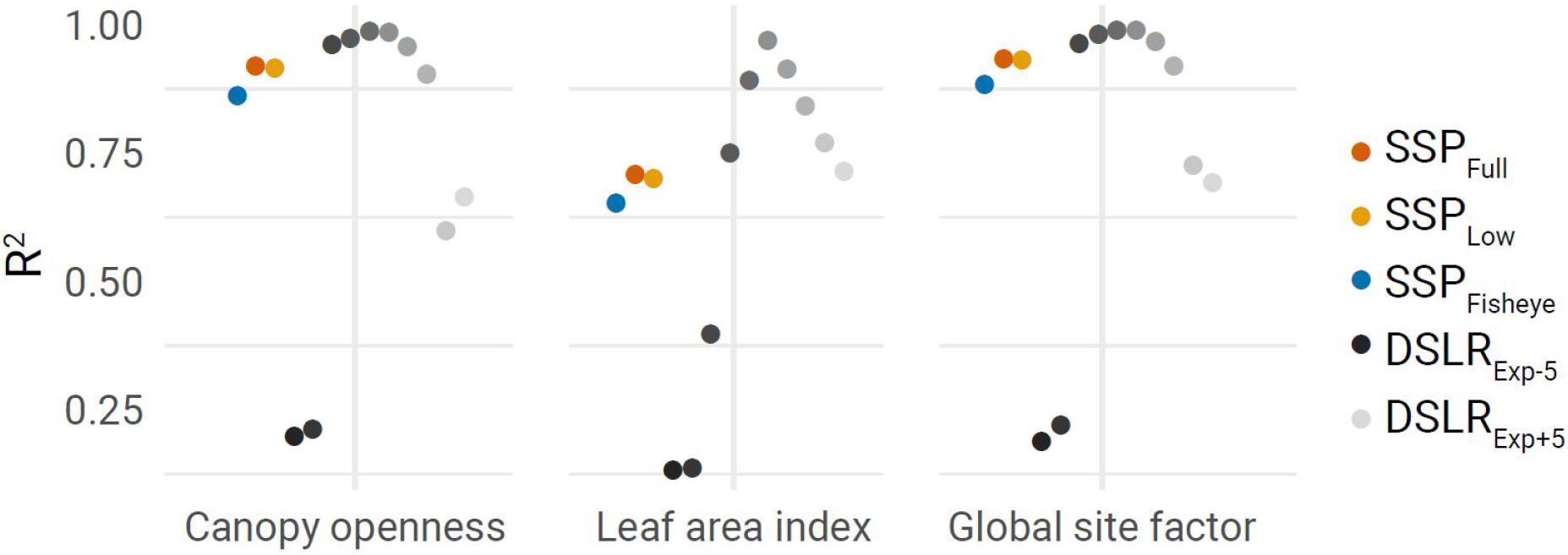
Correlation coefficients from independent OLS regression models predicting canopy structure and light environment values for reference (standard DSLR HP) from full resolution SSP HP (dark orange), low resolution SSP HP downsample to match the standard DSLR resolution (light orange), fisheye HP (blue), and DSLR HP with exposure adjusted from +5 to −5 (light to dark).

### 4.4 Comparison to 150° FOV fisheye HP

HP images emulating perpendicular 150° FOV fisheye images were moderately to highly correlated with the reference DSLR HP images for CO (R^2^ = 0.87), GSF (R^2^ = 0.89), and LAI (R^2^ = 0.47) but less so than true HP produced by spherical panoramas (Fig. 4, Table S3). Fisheye HP image values tended to result in overestimates of CO (+78%) and GSF (+73%) more than circular SSP HP (Table S3). However, the restricted FOV of the fisheye images were slightly more accurate in estimating LAI with respect to the reference images (−17%) compared to SSP HP (Table S3). Estimates from the fisheye HP method are comparable to DSLR HP images 1-2 stops overexposed.

## 5. Discussion

Smartphones have become nearly ubiquitous, yet researchers typically do not exploit even a fraction of their potential as a research tool. HP generated from smartphone spherical panoramas offers a highly accurate alternative to traditional DSLR HP with over 90% correlation with traditional methods for common canopy and light measures. The difference between estimates of canopy structure and light environment from spherical panoramas vary from the reference photos by about the same as over- or under-exposing images by 1-1.5 stops.

The primary differences between SSP HP and DSLR HP are that the former produces larger images, sharper resolution, and more even tonal range across the image. SSP HP generated with the Google Pixel 4a smartphone are over nine times larger than those taken with a DSLR. However, the simple increase in resolution does not account for the difference in clarity, as downsampled SSP HP images still retain more fine structure than DSLR HP images of the same size. When comparing DSLR HP images to those from SSP HP at the same resolution, it is clear that DSLR HP tends to underestimate the number of canopy gaps. This is due to low clarity causing adjacent pixels to bleed into each other, even when generated with industry standard lens and camera. Thus, the smallest gaps tend to be lost, as evidenced by higher relative gap size across all canopy densities, and small gaps tend to collapse into larger gaps more quickly. This second point can be seen in the way relative gap size increases with canopy openness much faster for DSLR HP images. Thus, although SSP HP images contain many more total gaps, the total gap area is similar, albeit with slightly more relative gap area as a consequence of retaining small gaps.

The difference in clarity is most likely a product of the restricted FOV of individual photos included in the panorama, improved sharpening through computational photography, and homogenous tone across zenith regions. In contrast, DSLR HP images suffer from glare and hazing associated with extremely wide FOV lenses. In addition, DSLR cameras struggle to evenly expose the entire hemisphere in a single exposure. For these reasons, SSP HP is likely to be far more robust to non-optimal lighting conditions, but more studies in variable skies are needed. An additional advantage of SSP HP is that, unlike DSLR methods, automatic exposure aided by HDR, effectively obviates the need for tedious manual exposure settings. Although this can introduce errors in binarization, minimal contrast stretching appears to solve the issue. Improvements in pixel classification beyond simple thresholding (e.g. Díaz, Negri and Lencinas, 2021) are likely to make this a nonissue even in direct sunlight conditions.

Artifacts generated by the panorama stitching process did not have a noticeable effect on canopy and light estimates but are a potential source of error. Care during image capture can reduce most cases of incongruity, however. Spherical panorama software assumes that all images are captured by rotating the camera around a single point in space (Fangi and Nardinocchi, 2013). When taking images by hand, it is easy to shift the phone, and therefore the image plane, during rotation. Practice in steady positioning helps. Also, the ability to immediately review panoramas on one’s screen phone or with stereoptic headsets lets researchers catch errors and retake panos in the field. This problem will attenuate as smartphone stitching software continues to improve (Luhmann, 2004).

Smartphone spherical panorama HP offers practical benefits over other methods in requiring no additional equipment other than a smartphone. Leveling is achieved automatically via the phone’s gyroscope. Geographic coordinates, elevation, and orientation are retained in the image metadata and could easily be integrated into an analysis pipeline. The fact that waterproof housings are cheap and readily available for smartphones is another benefit to field work, not to be overlooked. Coupled with the fact that SSP HP requires no tedious exposure settings, this method is highly amenable to citizen science projects with untrained data collectors. Although there is likely to be variation between smartphones which must be validated in future studies, the lack of lenses with idiosyncratic projections removes a major source of variability. All SSP HP images have polar projection by virtue of originating as spherical images.

The method presented here outperforms other methods of capturing HP with smartphones via clip-on fisheye lenses. Simulating Bianchi et al.’s (2017) method showed that the loss of information from restricting images to 150° FOV results in skewed but still relatively accurate estimates of canopy structure and light environment. However, the comparison is generous in that the simulated fisheye HP suffered no effects of poor optical quality associated with the small size of clip-on smartphone lenses.

Only the upper hemisphere was extracted from the spherical panoramas in this study, but other portions of the panorama could be extracted for other purposes. For instance, horizontal panorama could be used for estimating basal area (Fastie, 2010) or mapping stands (Lu *et al.*, 2019). The lower hemisphere could be useful in monitoring understory plants or ground cover composition. Researchers can even enter the spherical image with a virtual reality headset to identify species after leaving the field.

SSP HP solves many of the problems associated with traditional HP while offering many practical benefits for field applications. The ubiquity of smartphones and their ever-improving quality of software and optical hardware will only widen the range of spherical panoramic imagery applications in silviculture and forest ecology into the future.

## Acknowledgements

I would like to thank Vince Mow for access to Rockstock property and Yale University for access to Yale Preserve. I thank Dr. David Skelly, Logan Billet, Dahn-Young Dong, and Dr. Marlyse Duguid for providing thoughts on the methods and manuscript.

Funding for this project was provided by Yale Institute for Biospheric Studies.

## Supplementary materials

The following supplementary material is available: additional data summary tables, a diagram detailing the workflow used in this study, R code used to conduct statistical analysis and generate figures, AutoHotKey macro script to automate Gap Light Analyzer software, and BIMP plug-in recipe code for batch processing spherical panoramas in GIMP. Due to file size, image files are available from the author upon request.

## Data Availability

Code for analysis and image processing is available in the supplemental materials. Due to the large file sizes, raw images are available upon request to the author.

## Notes

### Competing Interest Statement

The authors have declared no competing interest.

### Summary of Updates

Minor typos corrections and updating the supplemental materials to include code.

## References

Anderson, M. C. (1964) ‘Studies of the woodland light climate I. The photographic computation of light condition’, Journal of Ecology, 52, pp. 27–41.

Barbero-García, I. et al. (2018) ‘Smartphone-based close-range photogrammetric assessment of spherical objects’, Photogrammetric Record, The, 33(162), pp. 283–299.

Beckschäfer, P. et al. (2013) ‘On the exposure of hemispherical photographs in forests’, iForest, 6(4), pp. 228–237.

Beckschäfer, P. (2015) Hemispherical_2.0: Batch processing hemispherical and canopy photographs with ImageJ - User manual. doi: 10.13140/RG.2.1.3059.4088.

Bianchi, S. et al. (2017) ‘Rapid assessment of forest canopy and light regime using smartphone hemispherical photography’, Ecology and evolution, 7(24), pp. 10556–10566.

Brown, M. and Lowe, D. G. (2007) ‘Automatic panoramic image stitching using invariant features’, International journal of computer vision, 74(1), pp. 59–73.

Brown, P. L., Doley, D. and Keenan, R. J. (2000) ‘Estimating tree crown dimensions using digital analysis of vertical photographs’, Agricultural and Forest Meteorology, 100(2), pp. 199–212.

Chianucci, F. (2020) ‘An overview of in situ digital canopy photography in forestry’, Canadian journal of forest research. Journal canadien de la recherche forestiere, 50(3), pp. 227–242.

Chianucci, F. and Cutini, A. (2012) ‘Digital hemispherical photography for estimating forest canopy properties: Current controversies and opportunities’, iForest - Biogeosciences and Forestry, 5(6). doi: 10.3832/ifor0775-005.

Díaz, G. M., Negri, P. A. and Lencinas, J. D. (2021) ‘Toward making canopy hemispherical photography independent of illumination conditions: A deep-learning-based approach’, Agricultural and Forest Meteorology, 296, p. 108234.

Fangi, G. and Nardinocchi, C. (2013) ‘Photogrammetric processing of spherical panoramas’, Photogrammetric Record, The, 28(143), pp. 293–311.

Fastie, C. L. (2010) ‘Estimating stand basal area from forest panoramas’, in Proceedings of the Fine International Conference on Gigapixel Imaging for Science. Fine International Conference on Gigapixel Imagingfor Science, Carnegie Mellon University, pp. 1–7.

Frazer, G. W., Canham, C. D. and Lertzman, K. P. (1999) Gap Light Analyzer (GLA): Imaging software to extract canopy structure and gap light transmission indices from true-color fisheye photographs, users manual and documentation. Burnaby, British Columbia: Simon Fraser University. Available at: http://rem-main.rem.sfu.ca/downloads/Forestry/GLAV2UsersManual.pdf.

Frazer, G. W., Trofymow, J. A. and Lertzman, K. P. (1997) A method for estimating canopy openness, effective leaf area index, and photosynthetically active photon flux density using hemispherical photography and computerized image analysis techniques. BC-X-373. Pacific Forestry Centre. Available at: http://citeseerx.ist.psu.edu/viewdoc/download?doi=10.1.1.477.483&rep=rep1&type=pdf.

Glatthorn, J. and Beckschäfer, P. (2014) ‘Standardizing the protocol for hemispherical photographs: accuracy assessment of binarization algorithms’, PloS one, 9(11), p. e111924.

Gonsamo, A., D’odorico, P. and Pellikka, P. (2013) ‘Measuring fractional forest canopy element cover and openness - definitions and methodologies revisited’, Oikos, 122(9), pp. 1283–1291.

Gunturk, B. K. (2017) ‘Super-resolution imaging’, in Lukac, R. (ed.) Computational Photography: Methods and Applications. CRC Press, pp. 175–208.

Halverson, M. A. et al. (2003) ‘Forest mediated light regime linked to amphibian distribution and performance’, Oecologia, 134(3), pp. 360–364.

Ignatov, A. et al. (2017) ‘DSLR-quality photos on mobile devices with deep convolutional networks’, arXiv [cs.CV]. Available at: http://arxiv.org/abs/1704.02470.

Jennings, S. B., Brown, N. D. and Sheil, D. (1999) ‘Assessing forest canopies and understorey illumination: canopy closure, canopy cover and other measures’, Forestry, 72(1), pp. 59–74.

Li, X. and Ratti, C. (2019) ‘Mapping the spatio-temporal distribution of solar radiation within street canyons of Boston using Google Street View panoramas and building height model’, Landscape and urban planning, 191, p. 103387.

Luhmann, T. (2004) ‘A historical review on panorama photogrammetry’, International Archives of the Photogrammetry, Remote Sensing and Spatial Information Sciences, 34(5/W16), p. 8.

Lukac, R. (2017) Computational Photography: Methods and Applications. CRC Press.

Lu, M.-K. et al. (2019) ‘Close-range photogrammetry with spherical panoramas for mapping spatial location and measuring diameters of trees under forest canopies’, Canadian journal of forest research. Journal canadien de la recherche forestiere, 49(8), pp. 865–874.

Macfarlane, C. (2011) ‘Classification method of mixed pixels does not affect canopy metrics from digital images of forest overstorey’, Agricultural and Forest Meteorology, 151(7), pp. 833–840.

Macfarlane, C. et al. (2014) ‘Digital canopy photography: Exposed and in the raw’, Agricultural and Forest Meteorology, 197, pp. 244–253.

Origo, N. et al. (2017) ‘Influence of levelling technique on the retrieval of canopy structural parameters from digital hemispherical photography’, Agricultural and Forest Meteorology, 237-238, pp. 143–149.

Prewitt, J. M. S. and Mendelsohn, M. L. (1966) ‘The analysis of cell images’, Annals of the New York Academy of Sciences, 128(3), pp. 1035–1053.

Promis, A. (2013) ‘Measuring and estimating the below-canopy light environment in a forest. a review’, 19(1)), pp. 139–146.

R Core Team (2019) R: A Language and Environment for Statistical Computing. Vienna, Austria: R Foundation for Statistical Computing. Available at: https://www.R-project.org/.

Rich, P. M. (1990) ‘Characterizing plant canopies with hemispherical photographs’, Remote Sensing Reviews, 5(1), pp. 13–29.

Szeliski, R. and Shum, H.-Y. (1997) ‘Creating full view panoramic image mosaics and environment maps’, in Proceedings of the 24th annual conference on Computer graphics and interactive techniques. USA: ACM Press/Addison-Wesley Publishing Co. (SIGGRAPH’97), pp. 251–258.

Tichý, L. (2016) ‘Field test of canopy cover estimation by hemispherical photographs taken with a smartphone’, Journal of vegetation science: official organ of the International Association for Vegetation Science. Edited by B. Collins, 27(2), pp. 427–435.

Zhang, Y., Chen, J. M. and Miller, J. R. (2005) ‘Determining digital hemispherical photograph exposure for leaf area index estimation’, Agricultural and Forest Meteorology, 133(1), pp. 166–181.

