## Supplementary material for "Evaluation of hemispherical photos extracted from smartphone spherical panorama images to estimate canopy structure and forest light environment": Analysis.R

```
###
# This code is associated with Arietta, AZA. 2020. Evaluation of hemispherical photos extracted from smartphone spherical panorama images to estimate canopy structure and forest light environment. doi: https://doi.org/10.1101/2020.12.15.422956
# This script takes the dataframe output from the script "ReadData.R" to run statistical analysis and make figures.
###

# R version 3.6.2
library(tidyverse) # tidyverse_1.3.0
library(ggridges) # ggridges_0.5.2
library(cowplot) # cowplot_1.1.0

## Set up visual settings
ColorSet <- data.frame(Process = c("Pixel_Hemi_Full_0_0.05", "Pixel_Hemi_Low_0_0.05", "Pixel_Fisheye_Mid_0_0.05", "DSLR_Hemi_Full_-5_0", "DSLR_Hemi_Full_-4_0", "DSLR_Hemi_Full_-3_0", "DSLR_Hemi_Full_-2_0", "DSLR_Hemi_Full_-1_0", "DSLR_Hemi_Full_1_0", "DSLR_Hemi_Full_2_0", "DSLR_Hemi_Full_3_0", "DSLR_Hemi_Full_4_0", "DSLR_Hemi_Full_5_0"),
           colorset = c("#D55E00", "#E69F00", "#0072B2", "grey14", "grey21", "grey28", "grey35", "grey42", "grey56", "grey63", "grey70", "grey78", "grey85"))
theme_set(theme_minimal())

## Set up the data
Imgs <- readRDS("./data/processed/YP_RS.rds") %>% 
  mutate(Sign = ifelse(Exp < 0, "n", "p")) %>%
  mutate(Process = paste(Camera, Method, Resolution, Exp, Contrast, sep = "_"),
         Process2 = paste(Camera, Method, Resolution, Sign, Exp, Contrast, sep = "_")) %>%
  filter(Process != "DSLR_Fisheye_Full_0_0") %>% # Remove the reference and mistake fisheye comparisson.
  mutate(Pix = ifelse(Camera == "DSLR" & Resolution == "Full", ((2885/2)^2)*pi, ((8704/2)^2)*pi)) %>%
  mutate(Pix = ifelse(Camera == "Pixel" & Resolution == "Mid", ((6049/2)^2)*pi, Pix)) %>%
  mutate(Pix = ifelse(Camera == "Pixel" & Resolution == "Low", ((2885/2)^2)*pi, Pix)) %>%
  mutate(av_gap_size = gap_area/no_of_gaps, rel_gap_size = (Canopy_open_perc/no_of_gaps))

Imgs %>% group_by(Location) %>% tally()
Imgs %>% group_by(Location, Site) %>% tally() %>% tally()
Imgs %>% group_by(Location, Site) %>% tally() %>% print(n = 72)
Imgs %>% group_by(Process) %>% tally()

# Create a long dataset
Imgs_long <-
  Imgs %>% pivot_longer(
    cols = c(
      Canopy_open_perc,
      Site_open_perc,
      LAI_4ring,
      LAI_5ring,
      Trans_Dir,
      Trans_Dif,
      Trans_Tot,
      SF_Dir,
      SF_Dif,
      GSF,
      no_of_gaps,
      gap_area,
      gap_fraction,
      av_gap_size,
      rel_gap_size
    ),
    names_to = "Measure",
    values_to = "Value"
  ) %>%
  mutate(Process = fct_relevel(Process, "Pixel_Hemi_Full_0_0.05", "Pixel_Hemi_Low_0_0.05", "Pixel_Fisheye_Mid_0_0.05", "DSLR_Hemi_Full_-5_0", "DSLR_Hemi_Full_-4_0", "DSLR_Hemi_Full_-3_0", "DSLR_Hemi_Full_-2_0", "DSLR_Hemi_Full_-1_0", "DSLR_Hemi_Full_1_0", "DSLR_Hemi_Full_2_0", "DSLR_Hemi_Full_3_0", "DSLR_Hemi_Full_4_0", "DSLR_Hemi_Full_5_0"))

unique(Imgs_long$Process)

Imgs_long <- Imgs_long %>%
  left_join(
    Imgs_long %>% filter(Process == "DSLR_Hemi_Full_0_0") %>% select(Site, Measure, Value) %>% rename(refValue = Value),
    by = c("Site", "Measure")
  ) %>%
  mutate(dif = Value - refValue) %>% # Join the reference dataset and compute a difference
  left_join(ColorSet, by = "Process") %>%
  mutate(Camera = fct_rev(Camera))

Imgs_long_proc <- Imgs_long %>% pivot_wider(id_cols = c(Site, Measure), names_from = Process, values_from = Value)

## Are there difference by location?
Loc.effect <- c()
for (i in unique(Imgs_long$Measure)) {
  mod <-lm(refValue ~ Location, data = Imgs_long %>% filter(Process == "DSLR_Hemi_Full_0_0" & Measure == i))
  summod <- summary(mod)
  DFnew <- data.frame(Measure = i, 
                      dif = round(summod$coefficients[2, 1], 2),
                      CI = paste0(round(summod$coefficients[2, 1] - summod$coefficients[2, 2]*1.96, 2), ", ", round(summod$coefficients[2, 1] + summod$coefficients[2, 2]*1.96, 2)),
                      p = round(summod$coefficients[2, 4], 2))
  Loc.effect <- bind_rows(Loc.effect, DFnew)
}
Loc.effect # The only significant difference between locations are LAI 5ring (but not 4ring) and number of gaps (but not gap area or gap fraction). So, I will ignore location.

# write.csv(Loc.effect %>% filter(Measure %in% c("Canopy_open_perc", "LAI_4ring", "GSF")), "./figs/Location_effect_Table.csv")

## Is it necessary to contrast-stretch smartphone photos for binarization?
Imgs_long %>%
  filter(Camera == "Pixel" & Method == "Hemi") %>%
  filter(Measure == "gap_fraction") %>%
  ggplot(aes(x = refValue, y = Value, col = as.factor(Contrast))) +
    geom_abline(slope = 1, intercept = 0, col = "grey", lty = 2) +
    geom_smooth(method = 'lm', se = F, size = 2) +
    geom_point(pch = 1) +
    coord_fixed(ratio = 1) +
    facet_grid(cols = vars(Resolution)) +
    theme(legend.position = 'none') +
    scale_color_manual(values = c("grey10", "grey60")) +
    labs(x = "Value (reference)") # There are definitely a few outliers with obviously misspecified thresholding without the contrast correction. So, I will include the contrast corrected versions from here on.
# ggsave("./figs/Smartphone_contrast.pdf", dpi = 300, width = 4, height = 5, units = "in")

Imgs_long <- Imgs_long %>% filter(Process != "Pixel_Hemi_Full_0_0" & Process != "Pixel_Hemi_Low_0_0" & Process != "DSLR_Hemi_Full_0_0") # Remove the smartphone image sets without contrast correction and remove the reference set since it is already included in the 'refValue'

## How do the smartphone processes compare to the fisheye method?

## Canopy gaps

Imgs_long %>%
  filter(Camera == "Pixel") %>%
  filter(Measure %in% unique(Imgs_long$Measure)[c(11, 12, 13, 15)]) %>%
  ggplot(aes(x = refValue, y = Value, col = colorset, fill = colorset)) +
    geom_abline(slope = 1, intercept = 0, col = "grey", lty = 2) +
    geom_smooth(method = 'lm', se = F) +
    geom_point(pch = 1) +
    facet_wrap(vars(Measure), scales = "free") +
    theme(legend.position = 'bottom', aspect.ratio = 1) +
    scale_color_identity() +
    theme(legend.position = "none") +
    labs(x = "Value (reference)")
  # Gap area is mainly a function of the total image area. 
  # Number of gaps is greater for all of the smartphone images and scales with resolution.
  # Gap fraction is basically identical for the circular images regardless of the resolution and very close to the reference with no skew. The fisheye method has a lower gap fraction estimate, which makes sense since it is excluding all of the gaps near the horizon. However, it is still pretty close.
# ggsave("./figs/Gap_smartphone.pdf", dpi = 300, width = 6.5, height = 6.5, units = "in")

Imgs_long %>%
  filter(Camera == "Pixel") %>%
  filter(Measure %in% unique(Imgs_long$Measure)[c(11)]) %>%
  ggplot(aes(x = refValue, y = Value, col = colorset, fill = colorset)) +
    geom_abline(slope = 1, intercept = 0, col = "grey", lty = 2) +
    geom_smooth(method = 'lm', se = F) +
    geom_point(pch = 1) +
    coord_fixed(ratio = 1, xlim = c(2250, 40000), ylim = c(2250, 40000)) +
    theme(legend.position = 'bottom') +
    labs(title = "Number of gaps") +
    scale_color_identity()
  # The number of gaps is much greater for the smartphone images, which seems due to better clarity since even at the same resolution, smartphone images have more gaps. It looks like smaller gaps aroung the horizon are mainly driving the trend because the number of gaps in the cell phone photos increases with the DSLR reference in the circular images, but not the fisheye images since the mask block those gaps at the horizon.

Imgs %>%
  filter(Process %in% c("Pixel_Hemi_Full_0_0.05", "Pixel_Hemi_Low_0_0.05", "Pixel_Fisheye_Mid_0_0.05", "DSLR_Hemi_Full_0_0")) %>%
  ggplot(aes(y = gap_fraction, x = no_of_gaps, col = Process, fill = Process)) +
    geom_smooth(method = 'lm', se = T) +
    geom_point(pch = 1) +
    theme(legend.position = 'bottom')
  # For the DLSR images, the gap fraction increases substantially with the number of gaps, but this is not the case for the smartphone images. This makes it seems like the increase in gaps in the smart phone images are due to the inclusion of more, smaller gaps.

Imgs_long %>%
  filter(Measure %in% unique(Imgs_long$Measure)[c(11)]) %>%
  ggplot(aes(x = refValue, y = Value, col = colorset, fill = colorset)) +
    geom_abline(slope = 1, intercept = 0, col = "grey", lty = 2) +
    geom_smooth(method = 'lm', se = F) +
    geom_point(pch = 1) +
    #coord_fixed(ratio = 1) +
    theme(legend.position = 'bottom') +
    labs(title = "Number of gaps") +
    facet_wrap(vars(Camera)) +
    scale_color_identity() +
    theme(legend.position = "none") +
    labs(x = "Number of gaps (reference)", y = "Number of gaps")
  # The number of gaps is not simply a matter of exposure. All of the DLSR exposure differences are lower than the smartphone images, with the exception of the the most overwxposed set.
# ggsave("./figs/GapNum_comparison.pdf", dpi = 300, width = 4, height = 4, units = "in")

Imgs_long %>%
  filter(Measure %in% unique(Imgs_long$Measure)[c(15)]) %>%
  ggplot(aes(x = refValue, y = Value, col = colorset, fill = colorset)) +
    geom_abline(slope = 1, intercept = 0, col = "grey", lty = 2) +
    geom_smooth(method = 'lm', se = F) +
    geom_point(pch = 1) +
    coord_fixed(ratio = 1) +
    theme(legend.position = 'bottom') +
    labs(title = "Relative gap size") +
    facet_wrap(vars(Camera)) +
    scale_color_identity() +
    theme(legend.position = "none") +
    coord_cartesian(ylim = c(0, 0.02)) +
    labs(x = "Relative gap size (reference)", y = "Relative gap size")
# ggsave("./figs/RelGapSize_comparison.pdf", dpi = 300, width = 4, height = 4, units = "in")

Imgs %>% 
  mutate(Camera = fct_rev(Camera)) %>%
  filter(Process != "Pixel_Hemi_Full_0_0" & Process != "Pixel_Hemi_Low_0_0" & Process != "DSLR_Hemi_Full_0_0") %>%
  left_join(ColorSet, by = "Process") %>%
  ggplot(aes(y = Canopy_open_perc, x = no_of_gaps, col = colorset, fill = colorset)) +
    geom_smooth(method = 'lm', se = F) +
    geom_point(pch = 1) +
    theme(legend.position = 'bottom') +
    facet_wrap(vars(Camera), scales = "free_x") +
    ylim(c(0,30)) +
    scale_color_identity() +
    scale_fill_identity() +
    labs(x = "Number of gaps", y = "CO")
  # However, it is clear that in the DSLR images, overexposure leads to blown out pixels that also increase the gap fraction, but this isn't the case for smartphone images. Note, here that I excluded points with gap fraction over 30% to reduce the influence of images that were misthresholded due to overexposure.
# ggsave("./figs/GapFrac_GapNum_comparisson.pdf", dpi = 300, width = 5, height = 3, units = "in")

Imgs %>% 
  filter(Process == "Pixel_Hemi_Low_0_0.05" | Process == "DSLR_Hemi_Full_0_0") %>%
  left_join(Imgs %>% filter(Process == "Pixel_Hemi_Full_0_0.05") %>% select(Site, no_of_gaps) %>% rename(Full_no_gaps = no_of_gaps), by = "Site") %>%
  ggplot(aes(y = rel_gap_size, x = Full_no_gaps, col = Process)) +
    geom_smooth(method = 'lm', se = F) +
    geom_point(pch = 1) +
    theme(legend.position = 'bottom') +
    coord_cartesian(ylim = c(0, 0.0125)) +
    scale_color_manual(values = c("grey50", "#D55E00", "#E69F00")) +
    labs(x = "Number of gaps", y = "Relative Gap Size")

Imgs %>% 
  mutate(Camera = fct_rev(Camera)) %>%
  filter(Process == "Pixel_Hemi_Low_0_0.05" | Process == "Pixel_Hemi_Full_0_0.05" | Process == "DSLR_Hemi_Full_0_0") %>%
  left_join(ColorSet, by = "Process") %>%
  mutate(CO_cat = cut(Canopy_open_perc, breaks = seq(0, 30, 5))) %>%
  filter(!is.na(CO_cat)) %>%
  ggplot(aes(y = rel_gap_size, x = Canopy_open_perc, col = Process, fill = Process, group = interaction(CO_cat, Process))) +
    geom_boxplot(alpha = 0.5) +
    labs(x = "CO", y = "Relative Gap Size") +
    scale_color_manual(values = c("grey50", "#D55E00", "#E69F00")) +
    scale_fill_manual(values = c("grey50", "#D55E00", "#E69F00")) +
    coord_cartesian(xlim = c(0, 15), ylim = c(0, 0.0035))
  # Higher resolution images have a smaller relative gap size. Even at the same res, smartphone images have a smaller rel gap size
  # Rel gap size increases with canopy openness, but faster for DSLR, probably because small gaps merge more.
# ggsave("./figs/RelGapSize_CO_comparisson.pdf", dpi = 300, width = 4, height = 2, units = "in")

Imgs %>% 
  mutate(Camera = fct_rev(Camera)) %>%
  filter(Process != "Pixel_Hemi_Full_0_0" & Process != "Pixel_Hemi_Low_0_0" & Process != "DSLR_Hemi_Full_0_0") %>%
  left_join(ColorSet, by = "Process") %>%
  ggplot(aes(y = rel_gap_size, x = no_of_gaps, col = colorset, fill = colorset)) +
    geom_smooth(method = 'lm', se = F) +
    #geom_point(pch = 1) +
    theme(legend.position = 'bottom') +
    facet_wrap(vars(Camera)) +
    coord_cartesian(ylim = c(0,0.012), xlim = c(0, 50)) +
    scale_color_identity() +
    scale_fill_identity() +
    labs(x = "Gap fraction", y = "Relative Gap Size")


## GLA estimates 

Imgs_long %>%
  filter(Camera == "Pixel") %>%
  filter(Measure %in% unique(Imgs_long$Measure)[c(1:10)]) %>%
  ggplot(aes(x = refValue, y = Value, col = colorset, fill = colorset)) +
    geom_abline(slope = 1, intercept = 0, col = "grey", lty = 2) +
    geom_smooth(method = 'lm', se = F) +
    geom_point(pch = 1) +
    facet_wrap(vars(Measure), scales = "free") +
    theme(legend.position = 'bottom', aspect.ratio = 1) +
    scale_color_identity() +
    scale_fill_identity() +
    labs(y = "Value", x = "Value (reference)")
# ggsave("./figs/GLAestimates_grid.pdf", dpi = 300, width = 6.5, height = 6, units = "in")

## LAI estimates

Imgs_long %>%
  filter(Camera == "Pixel") %>%
  filter(Measure %in% unique(Imgs_long$Measure)[c(3,4)]) %>%
  ggplot(aes(x = refValue, y = Value, col = colorset, fill = colorset)) +
    geom_abline(slope = 1, intercept = 0, col = "grey", lty = 2) +
    geom_smooth(method = 'lm', se = T) +
    geom_point(pch = 1) +
    facet_wrap(vars(Measure), scales = "free") +
    theme(legend.position = 'bottom', aspect.ratio = 1)  +
    scale_color_identity() +
    scale_fill_identity() +
    labs(x = "Value (reference)")
  # LAI shows the largest deviation. LAI 4ring is integrated over the 0-60 zenith where as LAI 5ring is integrated over 0-75. So the difference in fisheye to circular images is greater for LAI 5ring which includes more zentih area toward the horizon.
  # In general, it seems that DSLR underestimates small gaps, which inflates the LAI estimates, especially in dense canopies.
# ggsave("./figs/LAI_grid.pdf", dpi = 300, width = 6.5, height = 2.5, units = "in")

Imgs_long %>%
  filter(Measure %in% unique(Imgs_long$Measure)[c(3,4)]) %>%
  ggplot(aes(x = refValue, y = Value, col = colorset, fill = colorset)) +
    geom_abline(slope = 1, intercept = 0, col = "grey", lty = 2) +
    geom_smooth(method = 'lm', se = F) +
    geom_point(pch = 1) +
    facet_wrap(vars(Measure), scales = "free") +
    theme(legend.position = 'bottom', aspect.ratio = 1) +
    scale_color_identity() +
    scale_fill_identity()
  # LAI estimates are similar to DSLR images with 1-2 stops of overexposure.

## Canopy openness

Imgs_long %>%
  filter(Camera == "Pixel") %>%
  filter(Measure %in% unique(Imgs_long$Measure)[c(1:2)]) %>%
  ggplot(aes(x = refValue, y = Value, col = colorset, fill = colorset)) +
    geom_abline(slope = 1, intercept = 0, col = "grey", lty = 2) +
    geom_smooth(method = 'lm', se = T) +
    geom_point(pch = 1) +
    facet_wrap(vars(Measure), scales = "free") +
    theme(legend.position = 'bottom', aspect.ratio = 1) +
    scale_color_identity() +
    scale_fill_identity() +
    labs(x = "Value (reference)")
  # Smartphone images tend to exhibit greater openness in more dense canopies, probably because they include more small gaps.
# ggsave("./figs/Openness_grid.pdf", dpi = 300, width = 6.5, height = 2.5, units = "in")


## Site factors

Imgs_long %>%
  filter(Camera == "Pixel") %>%
  filter(Measure %in% unique(Imgs_long$Measure)[c(5:10)]) %>%
  ggplot(aes(x = refValue, y = Value, col = colorset, fill = colorset)) +
    geom_abline(slope = 1, intercept = 0, col = "grey", lty = 2) +
    geom_smooth(method = 'lm', se = T) +
    geom_point(pch = 1) +
    facet_wrap(vars(Measure), scales = "free") +
    theme(legend.position = 'bottom', aspect.ratio = 1) +
    scale_color_identity() +
    scale_fill_identity()  +
    labs(x = "Value (reference)")
  # Smartphone images also tend to exhibit greater SF values in denser canopies, likely also due to light transmitance through smaller gaps.
# ggsave("./figs/SiteFactor_grid.pdf", dpi = 300, width = 6.5, height = 5, units = "in")


Imgs_long %>%
  filter(Measure %in% unique(Imgs_long$Measure)[c(10)]) %>%
  ggplot(aes(x = refValue, y = Value, col = colorset, fill = colorset)) +
    geom_abline(slope = 1, intercept = 0, col = "grey", lty = 2) +
    geom_smooth(method = 'lm', se = F) +
    geom_point(pch = 1) +
    facet_wrap(vars(Measure), scales = "free") +
    theme(legend.position = 'bottom', aspect.ratio = 1) +
    scale_color_identity() +
    scale_fill_identity()
  # Deviance in GSF values are on par with over or under exposing by 1 stop.

## Create a table of estimates
Meas.deviance <- c()
for (i in unique(Imgs_long$Process)) {
  for (j in unique(Imgs_long$Measure)) {
    mod <-
      lm(refValue ~ Value, data = Imgs_long %>% filter(Process == i &
                                                       Measure == j))
    summod <- summary(mod)
    DFnew <- data.frame(
      Process = i,
      Measure = j,
      Intercept = summod$coefficients[1, 1],
      Int.SE = summod$coefficients[1,2],
      Int.p = summod$coefficients[1,4],
      Slope = summod$coefficients[2, 1],
      Slope.SE = summod$coefficients[2, 2],
      Slope.p = summod$coefficients[2, 4],
      R2 = summod$r.squared
    )
    Meas.deviance <- bind_rows(Meas.deviance, DFnew)
  }
}

Meas.deviance <- Meas.deviance %>% mutate(Process = fct_relevel(Process, c("Pixel_Hemi_Full_0_0.05", "Pixel_Hemi_Low_0_0.05", "Pixel_Fisheye_Mid_0_0.05", "DSLR_Hemi_Full_-5_0", "DSLR_Hemi_Full_-4_0", "DSLR_Hemi_Full_-3_0", "DSLR_Hemi_Full_-2_0", "DSLR_Hemi_Full_-1_0", "DSLR_Hemi_Full_1_0", "DSLR_Hemi_Full_2_0", "DSLR_Hemi_Full_3_0", "DSLR_Hemi_Full_4_0", "DSLR_Hemi_Full_5_0"))) %>%
  mutate(Process = fct_rev(Process)) %>%
  left_join(ColorSet, by = "Process") %>%
  left_join(Imgs %>% group_by(Process, Process2) %>% tally(), by = "Process") %>%
  separate(Process2, into = c("Camera", "Method", "Resolution", "Sign", "Exp", "Contrast")) %>%
  mutate(Exp = as.numeric(Exp)) %>%
  mutate(Exp = ifelse(Sign == "n", Exp * -1, Exp))

Meas.deviance %>% 
  filter(Measure != "Site_open_perc") %>%
  ggplot(aes(y = R2, x = Measure, col = colorset)) +
  geom_point(position = position_dodge(width = 0.5), size = 3, pch = 16) +
    scale_color_identity() +
    scale_fill_identity() +
  # coord_flip() +
  theme(panel.grid.major.y = element_line(color = "white"))

Meas.deviance %>% 
  filter(Measure != "Site_open_perc") %>%
  filter(Measure %in% unique(Imgs_long$Measure)[c(15, 12, 11, 1, 3, 10)]) %>%
  ggplot(aes(y = R2, x = Measure, col = colorset)) +
  geom_point(position = position_dodge(width = 0.5), size = 2, pch = 16) +
    scale_color_identity() +
    scale_fill_identity() +
  # coord_flip() +
  theme(panel.grid.major.y = element_line(color = "white"))
# ggsave("./figs/R2.pdf", dpi = 300, width = 6.5, height = 2, units = "in")

Meas.deviance %>% 
  filter(Measure != "Site_open_perc") %>%
  filter(Measure %in% unique(Imgs_long$Measure)[c(1, 3, 10)]) %>%
  ggplot(aes(y = R2, x = Measure, col = colorset)) +
  geom_point(position = position_dodge(width = 0.5), size = 3, pch = 16) +
    scale_color_identity() +
    scale_fill_identity() +
  # coord_flip() +
  theme(panel.grid.major.y = element_line(color = "white"))
# ggsave("./figs/R2.pdf", dpi = 300, width = 5.5, height = 2, units = "in")


Meas.deviance %>% filter(Measure == "rel_gap_size")

Meas.deviance %>% 
  mutate(Int.Lo = Intercept - (Int.SE*1.96), 
                         Int.Hi = Intercept + (Int.SE*1.96),
                         Slope.Lo = Slope - (Slope.SE*1.96),
                         Slope.Hi = Slope + (Slope.SE*1.96)) %>%
  mutate(Int.CI = paste0(round(Int.Lo, 2), ", ", round(Int.Hi, 2)),
         Slope.CI = paste0(round(Slope.Lo, 2), ", ", round(Slope.Hi, 2))) %>%
  mutate(R2 = round(R2, 2),
         Intercept = round(Intercept, 2),
         Slope = round(Slope, 2),
         Int.p = round(Int.p, 3),
         Slope.p = round(Slope.p, 3)) %>%
  select(Camera, Method, Resolution, Exp, Contrast, Measure, Intercept, Int.CI, Int.p, Slope, Slope.CI, Slope.p, R2) %>%
  mutate(Method = fct_rev(Method),
         Camera = fct_rev(Camera)) %>%
  arrange(Measure, Camera, Method, Resolution, Exp, Contrast) %>%
  filter(Camera == "Pixel")

Meas.deviance.table <- Meas.deviance %>% 
  filter(Measure %in% c("Canopy_open_perc", "LAI_4ring", "GSF")) %>%
  mutate(Int.Lo = Intercept - (Int.SE*1.96), 
                         Int.Hi = Intercept + (Int.SE*1.96),
                         Slope.Lo = Slope - (Slope.SE*1.96),
                         Slope.Hi = Slope + (Slope.SE*1.96)) %>%
  mutate(Int.CI = paste0(round(Int.Lo, 2), ", ", round(Int.Hi, 2)),
         Slope.CI = paste0(round(Slope.Lo, 2), ", ", round(Slope.Hi, 2))) %>%
  mutate(R2 = round(R2, 2),
         Intercept = round(Intercept, 2),
         Slope = round(Slope, 2),
         Int.p = round(Int.p, 3),
         Slope.p = round(Slope.p, 3)) %>%
  mutate(Intercept = paste0(Intercept, " (", Int.CI, "), ", Int.p),
         Slope = paste0(Slope, " (", Slope.CI, "), ", Slope.p)) %>%
  select(Camera, Method, Resolution, Exp, Contrast, Measure, Intercept, Slope, R2) %>%
  pivot_wider(id_cols = c(Camera, Method, Resolution, Exp, Contrast), names_from = Measure, values_from = c(Intercept, Slope, R2)) %>%
  mutate(Method = fct_rev(Method),
         Camera = fct_rev(Camera)) %>%
  arrange(Camera, Method, Resolution, Exp, Contrast)
# write.csv(Meas.deviance.table, "./figs/Measure_deviance_table_ALL.csv")
# write.csv(Meas.deviance.table, "./figs/Measure_deviance_table_LESS.csv")


Canopy_Comp_table <- Imgs_long %>%
  filter(Measure == "Canopy_open_perc") %>%
  group_by(Process, Measure) %>%
  mutate(perc.dif = (dif/refValue)*100) %>%
  summarise(Mean.ref = mean(refValue), Mean.val = mean(Value), Mean.dif = mean(dif), sd.dif = sd(dif), Med.dif = median(dif), Mean.perc.dif = mean(perc.dif), Med.perc.dif = median(perc.dif)) %>%
  mutate_at(vars(-"Measure", -"Process"), list(function(x)(round(x, 2)))) %>%
  mutate(Dif = paste0(Mean.dif, " (", sd.dif, ")")) %>%
  select(Process, Measure, Dif, Med.dif, Mean.perc.dif, Med.perc.dif) %>%
bind_rows(Imgs_long %>%
  filter(Measure == "LAI_5ring") %>%
  group_by(Process, Measure) %>%
  mutate(perc.dif = (dif/refValue)*100) %>%
  summarise(Mean.ref = mean(refValue), Mean.val = mean(Value), Mean.dif = mean(dif), sd.dif = sd(dif), Med.dif = median(dif), Mean.perc.dif = mean(perc.dif), Med.perc.dif = median(perc.dif)) %>%
  mutate_at(vars(-"Measure", -"Process"), list(function(x)(round(x, 2)))) %>%
  mutate(Dif = paste0(Mean.dif, " (", sd.dif, ")")) %>%
  select(Process, Measure, Dif, Med.dif, Mean.perc.dif, Med.perc.dif)) %>%
bind_rows(Imgs_long %>%
  filter(Measure == "GSF") %>%
  group_by(Process, Measure) %>%
  mutate(perc.dif = (dif/refValue)*100) %>%
  summarise(Mean.ref = mean(refValue), Mean.val = mean(Value), Mean.dif = mean(dif), sd.dif = sd(dif), Med.dif = median(dif), Mean.perc.dif = mean(perc.dif), Med.perc.dif = median(perc.dif)) %>%
  mutate_at(vars(-"Measure", -"Process"), list(function(x)(round(x, 2)))) %>%
  mutate(Dif = paste0(Mean.dif, " (", sd.dif, ")")) %>%
  select(Process, Measure, Dif, Med.dif, Mean.perc.dif, Med.perc.dif)) %>%
filter(Process != "DSLR_Hemi_Full_0_0" & Process != "Pixel_Hemi_Low_0_0" & Process != "Pixel_Hemi_Full_0_0")

# write.csv(Canopy_Comp_table, "./figs/Table_comp_canopy.csv")

Image_Comp_table <- Imgs_long %>%
  filter(Measure == "gap_area") %>%
  mutate_at(vars("Value", "refValue", "dif"), function(x)(x/1000000)) %>%
  group_by(Process, Measure) %>%
  mutate(perc.dif = (dif/refValue)*100) %>%
  summarise(Mean.ref = mean(refValue), Mean.val = mean(Value), Mean.dif = mean(dif), sd.dif = sd(dif), Med.dif = median(dif), Mean.perc.dif = mean(perc.dif), Med.perc.dif = median(perc.dif)) %>%
  mutate_at(vars(-"Measure", -"Process"), list(function(x)(round(x, 2)))) %>%
  mutate(Dif = paste0(Mean.dif, " (", sd.dif, ")")) %>%
  select(Process, Measure, Dif, Med.dif, Mean.perc.dif, Med.perc.dif) %>%
bind_rows(Imgs_long %>%
  filter(Measure == "no_of_gaps") %>%
  group_by(Process, Measure) %>%
  mutate(perc.dif = (dif/refValue)*100) %>%
  summarise(Mean.ref = mean(refValue), Mean.val = mean(Value), Mean.dif = mean(dif), sd.dif = sd(dif), Med.dif = median(dif), Mean.perc.dif = mean(perc.dif), Med.perc.dif = median(perc.dif)) %>%
  mutate_at(vars(-"Measure", -"Process"), list(function(x)(round(x, 2)))) %>%
  mutate(Dif = paste0(Mean.dif, " (", sd.dif, ")")) %>%
  select(Process, Measure, Dif, Med.dif, Mean.perc.dif, Med.perc.dif)) %>%
bind_rows(Imgs_long %>%
  filter(Measure == "rel_gap_size") %>%
  group_by(Process, Measure) %>%
  mutate(perc.dif = (dif/refValue)*100) %>%
  summarise(Mean.ref = mean(refValue), Mean.val = mean(Value), Mean.dif = mean(dif), sd.dif = sd(dif), Med.dif = median(dif), Mean.perc.dif = mean(perc.dif), Med.perc.dif = median(perc.dif)) %>%
  mutate_at(vars(-"Measure", -"Process"), list(function(x)(round(x, 2)))) %>%
  mutate(Dif = paste0(Mean.dif, " (", sd.dif, ")")) %>%
  select(Process, Measure, Dif, Med.dif, Mean.perc.dif, Med.perc.dif)) %>%
filter(Process != "DSLR_Hemi_Full_0_0" & Process != "Pixel_Hemi_Low_0_0" & Process != "Pixel_Hemi_Full_0_0")

# write.csv(Image_Comp_table, "./figs/Table_comp_images.csv")


Imgs_long %>%
  filter(Measure == "GSF") %>%
  group_by(Process, Measure) %>%
  mutate(perc.dif = (dif/refValue)*100) %>%
  summarise(Mean.dif = mean(dif), sd.dif = sd(dif), Med.dif = median(dif), Mean.perc.dif = mean(perc.dif), Med.perc.dif = median(perc.dif))

Imgs_long %>%
  filter(Measure == "no_of_gaps") %>%
  group_by(Process, Measure) %>%
  mutate(perc.dif = (dif/refValue)*100) %>%
  summarise(Mean.ref = mean(refValue), Mean.val = mean(Value), Mean.dif = mean(dif), sd.dif = sd(dif), Med.dif = median(dif), Mean.perc.dif = mean(perc.dif), Med.perc.dif = median(perc.dif))

Imgs_long %>%
  filter(Measure == "gap_area") %>%
  group_by(Process, Measure) %>%
  mutate(perc.dif = (dif/refValue)*100) %>%
  summarise(Mean.ref = mean(refValue), Mean.val = mean(Value), Mean.dif = mean(dif), sd.dif = sd(dif), Med.dif = median(dif), Mean.perc.dif = mean(perc.dif), Med.perc.dif = median(perc.dif))

Imgs_long %>%
  filter(Measure == "rel_gap_size") %>%
  group_by(Process, Measure) %>%
  mutate(perc.dif = (dif/refValue)*100) %>%
  summarise(Mean.ref = mean(refValue), Mean.val = mean(Value), Mean.dif = mean(dif), sd.dif = sd(dif), Med.dif = median(dif), Mean.perc.dif = mean(perc.dif), Med.perc.dif = median(perc.dif))

Imgs_long %>%
  filter(Measure == "LAI_5ring") %>%
  group_by(Process, Measure) %>%
  mutate(perc.dif = (dif/refValue)*100) %>%
  summarise(Mean.ref = mean(refValue), Mean.val = mean(Value), Mean.dif = mean(dif), sd.dif = sd(dif), Med.dif = median(dif), Mean.perc.dif = mean(perc.dif), Med.perc.dif = median(perc.dif))


Imgs_long %>%
  group_by(Measure) %>%
  summarise(min(refValue), max(refValue), mean(refValue), sd(refValue), median(refValue))

Imgs_long %>%
  group_by(Measure) %>%
  summarise(q25 = quantile(refValue, c(0.25)), q75 = quantile(refValue, c(0.75)))

Res_comparisson_table <- Imgs_long %>%
  filter(Process %in% c("Pixel_Hemi_Full_0_0.05", "Pixel_Hemi_Low_0_0.05")) %>%
  group_by(Measure, Process) %>%
  summarise(mean = mean(Value), sd = sd(Value)) %>%
  mutate(Process = ifelse(Process == "Pixel_Hemi_Full_0_0.05", "F", "L")) %>%
  pivot_wider(id_cols = c(Measure), names_from = Process, values_from = c(mean, sd)) %>%
  mutate(mean.dif.perc = ((mean_L - mean_F)/mean_F)*100, sd.dif.perc = ((sd_L - sd_F)/sd_F)*100) %>%
  filter(Measure %in% c("no_of_gaps", "rel_gap_size", "Canopy_open_perc", "LAI_5ring", "GSF", "gap_area")) %>%
  group_by(Measure) %>%
  mutate_at(vars(-"Measure"), list(function(x)(round(x, 2)))) %>%
  mutate(Full = paste0(mean_F, " (", sd_F, ")"), 
         Low = paste0(mean_L, " (", sd_L, ")")) %>%
  select(Measure, Full, Low, mean.dif.perc, sd.dif.perc)
# write.csv(Res_comparisson_table, "./figs/Res_Comp_Table.csv")


# Make ridgepot of deviation from the DSLR standard reference set
SF_dir <- Imgs_long %>%
  filter(Measure %in% unique(Imgs_long$Measure)[c(8)]) %>%
  mutate(Process = fct_rev(Process)) %>%
  ggplot(aes(x = dif, y = Process, fill = colorset)) +
  geom_vline(xintercept = 0) +
  geom_density_ridges(rel_min_height = 0.001) +
  facet_grid(cols = vars(Measure)) +
  coord_cartesian(xlim = c(-35, 40)) +
  scale_fill_identity() +
  theme(
    panel.grid.major.x = element_blank(),
    panel.grid.minor.x = element_blank(),
    axis.text.y = element_blank(),
    axis.title = element_blank()
  )

SF_dif <- Imgs_long %>%
    filter(Measure %in% unique(Imgs_long$Measure)[c(9)]) %>%
    mutate(Process = fct_rev(Process)) %>%
    ggplot(aes(
      x = dif, y = Process, fill = colorset
    )) +
    geom_vline(xintercept = 0) +
    geom_density_ridges(rel_min_height = 0.001) +
    facet_grid(cols = vars(Measure)) +
    coord_cartesian(xlim = c(-35, 40)) +
    scale_fill_identity() +
    theme(
      panel.grid.major.x = element_blank(),
      panel.grid.minor.x = element_blank(),
      axis.text.y = element_blank(),
      axis.title = element_blank()
    )

GSF <- Imgs_long %>%
    filter(Measure %in% unique(Imgs_long$Measure)[c(10)]) %>%
    mutate(Process = fct_rev(Process)) %>%
    ggplot(aes(
      x = dif, y = Process, fill = colorset
    )) +
    geom_vline(xintercept = 0) +
    geom_density_ridges(rel_min_height = 0.001) +
    facet_grid(cols = vars(Measure)) +
    coord_cartesian(xlim = c(-35, 40)) +
    scale_fill_identity() +
    theme(
      panel.grid.major.x = element_blank(),
      panel.grid.minor.x = element_blank(),
      axis.text.y = element_blank(),
      axis.title = element_blank()
    )

Trans_dir <- Imgs_long %>%
    filter(Measure %in% unique(Imgs_long$Measure)[c(5)]) %>%
    mutate(Process = fct_rev(Process)) %>%
    ggplot(aes(
      x = dif, y = Process, fill = colorset
    )) +
    geom_vline(xintercept = 0) +
    geom_density_ridges(rel_min_height = 0.001) +
    facet_grid(cols = vars(Measure)) +
    coord_cartesian(xlim = c(-10, 10)) +
    scale_fill_identity() +
    theme(
      panel.grid.major.x = element_blank(),
      panel.grid.minor.x = element_blank(),
      axis.text.y = element_blank(),
      axis.title = element_blank()
    )

Trans_dif <- Imgs_long %>%
    filter(Measure %in% unique(Imgs_long$Measure)[c(6)]) %>%
    mutate(Process = fct_rev(Process)) %>%
    ggplot(aes(
      x = dif, y = Process, fill = colorset
    )) +
    geom_vline(xintercept = 0) +
    geom_density_ridges(rel_min_height = 0.001) +
    facet_grid(cols = vars(Measure)) +
    coord_cartesian(xlim = c(-10, 10)) +
    scale_fill_identity() +
    theme(
      panel.grid.major.x = element_blank(),
      panel.grid.minor.x = element_blank(),
      axis.text.y = element_blank(),
      axis.title = element_blank()
    )

Trans_tot <- Imgs_long %>%
    filter(Measure %in% unique(Imgs_long$Measure)[c(7)]) %>%
    mutate(Process = fct_rev(Process)) %>%
    ggplot(aes(
      x = dif, y = Process, fill = colorset
    )) +
    geom_vline(xintercept = 0) +
    geom_density_ridges(rel_min_height = 0.001) +
    facet_grid(cols = vars(Measure)) +
    coord_cartesian(xlim = c(-10, 10)) +
    scale_fill_identity() +
    theme(
      panel.grid.major.x = element_blank(),
      panel.grid.minor.x = element_blank(),
      axis.text.y = element_blank(),
      axis.title = element_blank()
    )

CO <- Imgs_long %>%
    filter(Measure %in% unique(Imgs_long$Measure)[c(1)]) %>%
    mutate(Process = fct_rev(Process)) %>%
    ggplot(aes(
      x = dif, y = Process, fill = colorset
    )) +
    geom_vline(xintercept = 0) +
    geom_density_ridges(rel_min_height = 0.001) +
    facet_grid(cols = vars(Measure)) +
    coord_cartesian(xlim = c(-22, 22)) +
    scale_fill_identity() +
    theme(
      panel.grid.major.x = element_blank(),
      panel.grid.minor.x = element_blank(),
      axis.text.y = element_blank(),
      axis.title = element_blank()
    )

SO <- Imgs_long %>%
    filter(Measure %in% unique(Imgs_long$Measure)[c(2)]) %>%
    mutate(Process = fct_rev(Process)) %>%
    ggplot(aes(
      x = dif, y = Process, fill = colorset
    )) +
    geom_vline(xintercept = 0) +
    geom_density_ridges(rel_min_height = 0.001) +
    facet_grid(cols = vars(Measure)) +
    coord_cartesian(xlim = c(-22, 22)) +
    scale_fill_identity() +
    theme(
      panel.grid.major.x = element_blank(),
      panel.grid.minor.x = element_blank(),
      axis.text.y = element_blank(),
      axis.title = element_blank()
    )

LAI4 <- Imgs_long %>%
    filter(Measure %in% unique(Imgs_long$Measure)[3]) %>%
    mutate(Process = fct_rev(Process)) %>%
    ggplot(aes(
      x = dif, y = Process, fill = colorset
    )) +
    geom_vline(xintercept = 0) +
    geom_density_ridges(rel_min_height = 0.001) +
    facet_grid(cols = vars(Measure)) +
    #coord_cartesian(xlim = c(-18, 18)) +
    scale_fill_identity() +
    theme(
      panel.grid.major.x = element_blank(),
      panel.grid.minor.x = element_blank(),
      axis.text.y = element_blank(),
      axis.title = element_blank()
    )

LAI <- Imgs_long %>%
    filter(Measure %in% unique(Imgs_long$Measure)[3]) %>%
    mutate(Process = fct_rev(Process)) %>%
    ggplot(aes(
      x = dif, y = Process, fill = colorset
    )) +
    geom_vline(xintercept = 0) +
    geom_density_ridges(rel_min_height = 0.001) +
    facet_grid(cols = vars(Measure)) +
    #coord_cartesian(xlim = c(-15, 15)) +
    scale_fill_identity() +
    theme(
      panel.grid.major.x = element_blank(),
      panel.grid.minor.x = element_blank(),
      axis.text.y = element_blank(),
      axis.title = element_blank()
    )

Gap_num <- Imgs_long %>%
    filter(Measure %in% unique(Imgs_long$Measure)[11]) %>%
    mutate(Process = fct_rev(Process)) %>%
    ggplot(aes(
      x = dif, y = Process, fill = colorset
    )) +
    geom_vline(xintercept = 0) +
    geom_density_ridges(rel_min_height = 0.001) +
    facet_grid(cols = vars(Measure)) +
    #coord_cartesian(xlim = c(-40000, 40000)) +
    scale_fill_identity() +
    theme(
      panel.grid.major.x = element_blank(),
      panel.grid.minor.x = element_blank(),
      axis.text.y = element_blank(),
      axis.title = element_blank()
    )

Gap_area <- Imgs_long %>%
    filter(Measure %in% unique(Imgs_long$Measure)[12]) %>%
    mutate(Process = fct_rev(Process)) %>%
    ggplot(aes(
      x = dif/1000, y = Process, fill = colorset
    )) +
    geom_vline(xintercept = 0) +
    geom_density_ridges(rel_min_height = 0.001) +
    facet_grid(cols = vars(Measure)) +
    coord_cartesian(xlim = c(-2000, 8000)) +
    scale_fill_identity() +
    theme(
      panel.grid.major.x = element_blank(),
      panel.grid.minor.x = element_blank(),
      axis.text.y = element_blank(),
      axis.title = element_blank()
    )

Gap_frac <- Imgs_long %>%
    filter(Measure %in% unique(Imgs_long$Measure)[13]) %>%
    mutate(Process = fct_rev(Process)) %>%
    ggplot(aes(
      x = dif, y = Process, fill = colorset
    )) +
    geom_vline(xintercept = 0) +
    geom_density_ridges(rel_min_height = 0.001) +
    facet_grid(cols = vars(Measure)) +
    coord_cartesian(xlim = c(-22, 22)) +
    scale_fill_identity() +
    theme(
      panel.grid.major.x = element_blank(),
      panel.grid.minor.x = element_blank(),
      axis.text.y = element_blank(),
      axis.title = element_blank()
    )

Rel_gap_size <- Imgs_long %>%
    filter(Measure %in% unique(Imgs_long$Measure)[15]) %>%
    mutate(Process = fct_rev(Process)) %>%
    ggplot(aes(
      x = dif, y = Process, fill = colorset
    )) +
    geom_vline(xintercept = 0) +
    geom_density_ridges(rel_min_height = 0.001) +
    facet_grid(cols = vars(Measure)) +
    coord_cartesian(xlim = c(-0.003, 0.003)) +
    scale_fill_identity() +
    theme(
      panel.grid.major.x = element_blank(),
      panel.grid.minor.x = element_blank(),
      axis.text.y = element_blank(),
      axis.title = element_blank()
    )

plot_grid(CO, LAI4, GSF, Gap_area, Gap_num, Rel_gap_size)
# ggsave("./figs/DensRidges.pdf", dpi = 300, width = 6.5, height = 4.5, units = "in")
```
