## Supplementary material for "Evaluation of hemispherical photos extracted from smartphone spherical panorama images to estimate canopy structure and forest light environment": HistogramFigures.R

```
###
# This code is associated with Arietta, AZA. 2020. Evaluation of hemispherical photos extracted from smartphone spherical panorama images to estimate canopy structure and forest light environment. doi: https://doi.org/10.1101/2020.12.15.422956
# This script takes comparison images of site RS017 taken with DSLR and smartphone and outputs figures of the pixel value histograms.
###

# R version 3.6.2 (2019-12-12)
library(imager) # imager_0.42.3
library(tidyverse) # tidyverse_1.3.0
sessionInfo()

# Set up the visual environment
theme_set(theme_void())

# Create histograms of the two images and save them as pdfs
DSLR <- load.image('./data/raw/RS017_DSLRHP.png') %>% 
  as.data.frame() %>%
  filter(value > 0 & value < 1) %>%
  ggplot(aes(x = value*255)) +
  geom_histogram(binwidth = 1)

SSP <- load.image('./data/raw/RS017_SSPHP.png') %>% 
  as.data.frame() %>%
  filter(value > 0 & value < 1) %>%
  ggplot(aes(x = value*255)) +
  geom_histogram(binwidth = 1)

ggsave(DSLR, "./figs/HistogramDSLR.pdf", dpi = 300, width = 3, height = 1, units = "in")
ggsave(SSP, "./figs/HistogramSSP.pdf", dpi = 300, width = 3, height = 1, units = "in")
```
