## Supplementary material for "Evaluation of hemispherical photos extracted from smartphone spherical panorama images to estimate canopy structure and forest light environment": ReadDatafiles.R

```
###
# This code is associated with Arietta, AZA. 2020. Evaluation of hemispherical photos extracted from smartphone spherical panorama images to estimate canopy structure and forest light environment. doi: https://doi.org/10.1101/2020.12.15.422956
# This script takes the raw output files from Hemispherical 2.0 plugin for ImageJ and raw output from Gap Light Analyzer and compiles all into a single dataframe for further analysis.
###

library(tidyverse)

theme_set(theme_minimal())
# DSLR hemispherical lens

## Rockstock 2020 ====
## . Hemipix ====
## . . . DSLR ====
RS_Hpix_DSLR_main <- read_table2("./data/raw/RS2020_data/Exp0_hemipix.txt") %>%
  mutate(Exp = 0) %>%
  bind_rows(read_table2("./data/raw/RS2020_data/Minus1_hemipix.txt") %>%
    mutate(Exp = -1)) %>%
  bind_rows(read_table2("./data/raw/RS2020_data/Minus2_hemipix.txt") %>%
    mutate(Exp = -2)) %>%
  bind_rows(read_table2("./data/raw/RS2020_data/Minus3_hemipix.txt") %>%
    mutate(Exp = -3)) %>%
  bind_rows(read_table2("./data/raw/RS2020_data/Minus4_hemipix.txt") %>%
    mutate(Exp = -4)) %>%
  bind_rows(read_table2("./data/raw/RS2020_data/Minus5_hemipix.txt") %>%
    mutate(Exp = -5)) %>%
  bind_rows(read_table2("./data/raw/RS2020_data/Plus1_hemipix.txt") %>%
    mutate(Exp = 1)) %>%
  bind_rows(read_table2("./data/raw/RS2020_data/Plus2_hemipix.txt") %>%
    mutate(Exp = 2)) %>%
  bind_rows(read_table2("./data/raw/RS2020_data/Plus3_hemipix.txt") %>%
    mutate(Exp = 3)) %>%
  bind_rows(read_table2("./data/raw/RS2020_data/Plus4_hemipix.txt") %>%
    mutate(Exp = 4)) %>%
  bind_rows(read_table2("./data/raw/RS2020_data/Plus5_hemipix.txt") %>%
    mutate(Exp = 5)) %>%
  mutate(Camera = "DSLR", Resolution = "Full", Contrast = 0, Method = "Hemi") %>%
  filter(photo != "00AARegistration3.jpg") %>%
  separate(photo, into = c("Location", "Site", "Ext")) %>%
  select(-Ext) %>%
  filter(Site != "008", Site != "026")


RS_Hpix_DSLR_overlay <- read_table2("./data/raw/RS2020_data/Overlay_hemipix.txt") %>%
  mutate(Exp = 0, Camera = "DSLR", Resolution = "Full", Contrast = 0, Method = "Fisheye") %>%
  filter(photo != "00AARegistration3.jpg") %>%
  separate(photo, into = c("Location", "Site", "a", "b")) %>%
  select(-a, -b) %>%
  filter(Site != "008", Site != "026")

## . . . Pixel spheres ====
RS_Hpix_Pixel_full_contrast <- read_table2("./data/raw/RS2020_data/Polar_contrast05_hemipix_B1.txt") %>%
  mutate(Exp = 0, Contrast = 0.05, Resolution = "Full") %>%
  bind_rows(read_table2("./data/raw/RS2020_data/Polar_contrast05_hemipix_B2.txt") %>%
              mutate(Exp = 0, Contrast = 0.05, Resolution = "Full")) %>%
  separate(photo, into = c("Location", "a", "Site", "b", "c")) %>%
  select(-a, -b, -c) %>%
  mutate(Camera = "Pixel")

RS_Hpix_Pixel_full_original <- read_table2("./data/raw/RS2020_data/Polar_hemipix_B1.txt") %>%
  mutate(Exp = 0, Contrast = 0, Resolution = "Full") %>%
  bind_rows(read_table2("./data/raw/RS2020_data/Polar_hemipix_B12.txt") %>%
              mutate(Exp = 0, Contrast = 0, Resolution = "Full")) %>%
  separate(photo, into = c("Location", "a", "Site", "b")) %>%
  select(-a, -b) %>%
  mutate(Camera = "Pixel")

RS_Hpix_Pixel_low_original <- read_table2("./data/raw/RS2020_data/Polar_sm_hemipix.txt") %>%
  mutate(Exp = 0, Contrast = 0, Resolution = "Low") %>%
  separate(photo, into = c("a", "Location", "b", "Site", "c")) %>%
  select(-a, -b, -c) %>%
  mutate(Camera = "Pixel")

RS_Hpix_Pixel_low_contrast <- read_table2("./data/raw/RS2020_data/Polar_sm_contrast05_hemipix.txt") %>%
  mutate(Exp = 0, Contrast = 0.05, Resolution = "Low") %>%
  separate(photo, into = c("a", "Location", "b", "Site", "c")) %>%
  select(-a, -b, -c) %>%
  mutate(Camera = "Pixel") 

RS_Hpix_Pixel_fisheye <- read_table2("./data/raw/RS2020_data/RS2020_Pixelfisheye_hpix.txt") %>%
  mutate(Exp = 0, Contrast = 0.05, Resolution = "Mid") %>%
  separate(photo, into = c("Location", "b", "Site", "c", "d")) %>%
  select(-b, -c, -d) %>%
  mutate(Camera = "Pixel", Method = "Fisheye") 

RS_Hpix_Pixel_full <- RS_Hpix_Pixel_full_original %>%
  bind_rows(RS_Hpix_Pixel_full_contrast) %>%
  bind_rows(RS_Hpix_Pixel_low_original) %>%
  bind_rows(RS_Hpix_Pixel_low_contrast) %>%
  mutate(Method = "Hemi") %>%
  bind_rows(RS_Hpix_Pixel_fisheye)

## . . Combined ====

RS_Hemipix_all <- bind_rows(RS_Hpix_DSLR_main,
                         RS_Hpix_DSLR_overlay,
                         RS_Hpix_Pixel_full)

RS_Hemipix_all %>% group_by(Camera, Exp, Resolution, Contrast, Method) %>% tally()


## . GLA ====
## . . . DSLR ====
RS_GLA_DSLR_main <- read_delim("./data/raw/RS2020_data/GLAestimates_2.sum", ";", escape_double = FALSE, trim_ws = TRUE) %>%
  rename(photo = 'User Field 1', 
         Canopy_open_perc = '% Cnpy Open',
         Site_open_perc = '% Site Open',
         LAI_4ring = 'LAI 4Ring',
         LAI_5ring = 'LAI 5Ring',
         Trans_Dir = 'Trans Dir',
         Trans_Dif = 'Trans Dif',
         Trans_Tot = 'Trans Tot',
         SF_Dir = '% Trans Dir',
         SF_Dif = '% Trans Dif',
         GSF = '% Trans Tot') %>%
  select(photo, Canopy_open_perc, Site_open_perc, LAI_4ring, LAI_5ring, Trans_Dir, Trans_Dif, Trans_Tot, SF_Dir, SF_Dif, GSF) %>% 
  filter(photo != "00AARegistration.bmp") %>%
  separate(photo, into = c("sign", "val", "Location", "Site", "T", "Ext")) %>%
  mutate(sign = ifelse(sign == "n", -1, 1), val = as.numeric(val)) %>%
  mutate(Exp = sign * val) %>%
  select(-sign, -val, -"T", -Ext) %>%
  mutate(Location = "Rockstock2020",
         Camera = "DSLR",
         Resolution = "Full",
         Contrast = 0,
         Method = "Hemi") %>%
  filter(Site != "008", Site != "026")

RS_GLA_DSLR_overlay <- read_delim("./data/raw/RS2020_data/GLA_DSLR_overlay150.sum", ";", escape_double = FALSE, trim_ws = TRUE) %>%
  rename(photo = 'User Field 1', 
         Canopy_open_perc = '% Cnpy Open',
         Site_open_perc = '% Site Open',
         LAI_4ring = 'LAI 4Ring',
         LAI_5ring = 'LAI 5Ring',
         Trans_Dir = 'Trans Dir',
         Trans_Dif = 'Trans Dif',
         Trans_Tot = 'Trans Tot',
         SF_Dir = '% Trans Dir',
         SF_Dif = '% Trans Dif',
         GSF = '% Trans Tot') %>%
  select(photo, Canopy_open_perc, Site_open_perc, LAI_4ring, LAI_5ring, Trans_Dir, Trans_Dif, Trans_Tot, SF_Dir, SF_Dif, GSF) %>% 
  filter(photo != "00AARegistration.bmp") %>%
  separate(photo, into = c("a", "b", "c", "Site", "Ext")) %>%
  select(-a, -b, -c, -Ext) %>%
  mutate(Location = "Rockstock2020",
         Camera = "DSLR",
         Resolution = "Full",
         Contrast = 0,
         Method = "Fisheye",
         Exp = 0) %>%
  filter(Site != "008", Site != "026")

## . . . Pixel spheres ====

RS_GLA_Pixel_full <- read_delim("./data/raw/RS2020_data/Polar_lg_and_contrast05_GLAoutput.sum", ";", escape_double = FALSE, trim_ws = TRUE) %>%
  rename(photo = 'User Field 1', 
         Canopy_open_perc = '% Cnpy Open',
         Site_open_perc = '% Site Open',
         LAI_4ring = 'LAI 4Ring',
         LAI_5ring = 'LAI 5Ring',
         Trans_Dir = 'Trans Dir',
         Trans_Dif = 'Trans Dif',
         Trans_Tot = 'Trans Tot',
         SF_Dir = '% Trans Dir',
         SF_Dif = '% Trans Dif',
         GSF = '% Trans Tot') %>%
  select(photo, Canopy_open_perc, Site_open_perc, LAI_4ring, LAI_5ring, Trans_Dir, Trans_Dif, Trans_Tot, SF_Dir, SF_Dif, GSF) %>% 
  filter(photo != "00_Registration.bmp") %>%
  separate(photo, into = c("a", "b", "Site", "Site2", "c")) %>%
  mutate(Contrast = ifelse(Site == "c05", 0.05, 0)) %>% 
  mutate(Site = ifelse(Contrast == 0, Site, Site2)) %>%
  select(-a, -b, -c, -Site2) %>%
  mutate(Location = "Rockstock2020",
         Camera = "Pixel",
         Resolution = "Full",
         Exp = 0,
         Method = "Hemi")

RS_GLA_Pixel_low_original <- read_delim("./data/raw/RS2020_data/Polar_sm_GLA.sum", ";", escape_double = FALSE, trim_ws = TRUE) %>%
  rename(photo = 'User Field 1', 
         Canopy_open_perc = '% Cnpy Open',
         Site_open_perc = '% Site Open',
         LAI_4ring = 'LAI 4Ring',
         LAI_5ring = 'LAI 5Ring',
         Trans_Dir = 'Trans Dir',
         Trans_Dif = 'Trans Dif',
         Trans_Tot = 'Trans Tot',
         SF_Dir = '% Trans Dir',
         SF_Dif = '% Trans Dif',
         GSF = '% Trans Tot') %>%
  select(photo, Canopy_open_perc, Site_open_perc, LAI_4ring, LAI_5ring, Trans_Dir, Trans_Dif, Trans_Tot, SF_Dir, SF_Dif, GSF) %>% 
  filter(photo != "00_Registration.bmp") %>%
  separate(photo, into = c("a", "b", "c", "Site", "d")) %>%
  mutate(Contrast = 0) %>%
  select(-a, -b, -c, -d) %>%
  mutate(Location = "Rockstock2020",
         Camera = "Pixel",
         Resolution = "Low",
         Exp = 0,
         Method = "Hemi")

RS_GLA_Pixel_low_contrast <- read_delim("./data/raw/RS2020_data/Polar_sm_contrast05_GLA.sum", ";", escape_double = FALSE, trim_ws = TRUE) %>%
  rename(photo = 'User Field 1', 
         Canopy_open_perc = '% Cnpy Open',
         Site_open_perc = '% Site Open',
         LAI_4ring = 'LAI 4Ring',
         LAI_5ring = 'LAI 5Ring',
         Trans_Dir = 'Trans Dir',
         Trans_Dif = 'Trans Dif',
         Trans_Tot = 'Trans Tot',
         SF_Dir = '% Trans Dir',
         SF_Dif = '% Trans Dif',
         GSF = '% Trans Tot') %>%
  select(photo, Canopy_open_perc, Site_open_perc, LAI_4ring, LAI_5ring, Trans_Dir, Trans_Dif, Trans_Tot, SF_Dir, SF_Dif, GSF) %>% 
  filter(photo != "00_Registration.bmp") %>%
  separate(photo, into = c("a", "b", "c", "d", "Site", "e")) %>%
  mutate(Contrast = 0.05) %>%
  select(-a, -b, -c, -d, -e) %>%
  mutate(Location = "Rockstock2020",
         Camera = "Pixel",
         Resolution = "Low",
         Exp = 0,
         Method = "Hemi")

RS_GLA_Pixel_fisheye <- read_delim("./data/raw/RS2020_data/RS2020_Pixelfisheye_GLA.sum", ";", escape_double = FALSE, trim_ws = TRUE) %>%
  rename(photo = 'User Field 1', 
         Canopy_open_perc = '% Cnpy Open',
         Site_open_perc = '% Site Open',
         LAI_4ring = 'LAI 4Ring',
         LAI_5ring = 'LAI 5Ring',
         Trans_Dir = 'Trans Dir',
         Trans_Dif = 'Trans Dif',
         Trans_Tot = 'Trans Tot',
         SF_Dir = '% Trans Dir',
         SF_Dif = '% Trans Dif',
         GSF = '% Trans Tot') %>%
  select(photo, Canopy_open_perc, Site_open_perc, LAI_4ring, LAI_5ring, Trans_Dir, Trans_Dif, Trans_Tot, SF_Dir, SF_Dif, GSF) %>%
  filter(photo != "00_Reg_6049.bmp") %>%
  separate(photo, into = c("a", "Site", "c", "Ext")) %>%
  select(-a, -c, -Ext) %>%
  mutate(Location = "Rockstock2020",
         Camera = "Pixel",
         Resolution = "Mid",
         Contrast = 0.05,
         Method = "Fisheye",
         Exp = 0)

# . . Combined ====
RS_GLA_all <- bind_rows(RS_GLA_DSLR_main,
                     RS_GLA_DSLR_overlay,
                     RS_GLA_Pixel_full,
                     RS_GLA_Pixel_low_original,
                     RS_GLA_Pixel_low_contrast,
                     RS_GLA_Pixel_fisheye)

RS_GLA_all %>% group_by(Camera, Exp, Resolution, Contrast, Method) %>% tally()

### . RS combined ====

Rockstock <- left_join(
  RS_GLA_all,
  RS_Hemipix_all %>% select(-Index),
  by = c("Camera", "Exp", "Resolution", "Contrast", "Location", "Site", "Method")
) %>%
  left_join(
    RS_GLA_all %>% filter(Camera == "DSLR" & Exp == 0 & Method == "Hemi") %>%
      mutate(stdGSF = GSF) %>%
      select(stdGSF, Site),
    by = c("Site")
  ) %>%
  mutate(Site = fct_reorder(Site, stdGSF)) %>%
  mutate(Site = paste0("RS", Site))

Rockstock %>%
  filter(Camera == "DSLR") %>%
  ggplot(aes(x = stdGSF, y = GSF, col = Exp)) +
  geom_point() +
  geom_point(data = Rockstock %>% filter(Camera == "DSLR" & Exp ==0), col = "firebrick") +
  scale_color_gradient(high = "grey80", low = "black") +
  coord_fixed(ratio = 1) +
  xlim(c(0, 100)) +
  ylim(c(0, 100))

Rockstock %>%
  filter(Camera == "Pixel") %>%
  ggplot(aes(x = stdGSF, y = GSF, col = Resolution)) +
  geom_abline(slope = 1, intercept = 0, col = "grey80") +
  geom_point() +
  coord_fixed(ratio = 1) +
  xlim(c(0, 100)) +
  ylim(c(0, 100)) +
  facet_wrap(vars(Contrast))


## Yale Preserve 2020 ====
## . Hemipix ====
## . . . DSLR ====

YP_Hpix_DSLR_main <- read_table2("./data/raw/YP2020_data/YP2020_DSLR_hpix_1.txt") %>%
  bind_rows(read_table2("./data/raw/YP2020_data/YP2020_DSLR_hpix_2.txt")) %>%
  bind_rows(read_table2("./data/raw/YP2020_data/YP2020_DSLR_hpix_3.txt")) %>%
  bind_rows(read_table2("./data/raw/YP2020_data/YP2020_DSLR_hpix_4.txt")) %>%
  mutate(Camera = "DSLR", Resolution = "Full", Contrast = 0, Method = "Hemi") %>%
  filter(photo != "00AARegistration3.jpg") %>%
  separate(photo, into = c("sign", "val", "Pond", "Site", "Ext")) %>%
  select(-Ext) %>%
  mutate(sign = ifelse(sign == "n", -1, 1)) %>%
  mutate(Exp = as.numeric(sign) * as.numeric(val)) %>%
  select(-sign, -val, -Index)

YP_Hpix_DSLR_overlay <- read_table2("./data/raw/YP2020_data/YP2020_DSLRfisheye_hpix.txt") %>%
  mutate(Exp = 0, Camera = "DSLR", Resolution = "Full", Contrast = 0, Method = "Fisheye") %>%
  filter(photo != "00AARegistration3.jpg") %>%
  separate(photo, into = c("sign", "val", "Pond", "Site", "Ext")) %>%
  select(-Ext) %>%
  mutate(sign = ifelse(sign == "n", -1, 1)) %>%
  mutate(Exp = as.numeric(sign) * as.numeric(val)) %>%
  select(-sign, -val, -Index)

## . . . Pixel spheres ====
YP_Hpix_Pixel_full <- read_table2("./data/raw/YP2020_data/YP2020_Pixel_c_hpix_1.txt") %>%
  bind_rows(read_table2("./data/raw/YP2020_data/YP2020_Pixel_c_hpix_2.txt")) %>%
  bind_rows(read_table2("./data/raw/YP2020_data/YP2020_Pixel_hpix_1.txt")) %>%
  bind_rows(read_table2("./data/raw/YP2020_data/YP2020_Pixel_hpix_2.txt")) %>%
  separate(photo, into = c("Pond", "Site", "Contrast", "Ext")) %>%
  mutate(Contrast = ifelse(Contrast == "c", 0.05, 0)) %>%
  mutate(Exp = 0, Resolution = "Full", Camera = "Pixel", Method = "Hemi") %>%
  select(-Ext, -Index)

YP_Hpix_Pixel_lo <- read_table2("./data/raw/YP2020_data/YP2020_Pixel_lo_hpix.txt") %>%
  separate(photo, into = c("Pond", "Site", "Contrast", "res", "Ext")) %>%
  mutate(Contrast = ifelse(Contrast == "c", 0.05, 0)) %>%
  mutate(Exp = 0, Resolution = "Low", Camera = "Pixel", Method = "Hemi") %>%
  select(-Ext, -res, -Index)

YP_Hpix_Pixel_fisheye <- read_table2("./data/raw/YP2020_data/YP2020_Pixelfisheye_hpix.txt") %>%
  mutate(Exp = 0, Camera = "Pixel", Resolution = "Mid", Contrast = 0.05, Method = "Fisheye") %>%
  separate(photo, into = c("Pond", "Site", "Ext")) %>%
  select(-Ext)

## . . Combined ====

YP_Hpix_all <- YP_Hpix_DSLR_main %>%
  bind_rows(YP_Hpix_DSLR_overlay,
            YP_Hpix_Pixel_full,
            YP_Hpix_Pixel_lo,
            YP_Hpix_Pixel_fisheye) %>%
  mutate(Site = paste0(Pond, "_", Site), Location = "YP2020") %>%
  select(-Pond)

YP_Hpix_all %>% group_by(Camera, Exp, Resolution, Contrast, Method) %>% tally()

YP_Hpix_all %>% filter(is.na(no_of_gaps))

## . GLA ====
## . . . DSLR ====

YP_GLA_DSLR_main <- read_delim("./data/raw/YP2020_data/YP2020_DSLR_GLA.sum", ";", escape_double = FALSE, trim_ws = TRUE) %>%
  rename(photo = 'User Field 1', 
         Canopy_open_perc = '% Cnpy Open',
         Site_open_perc = '% Site Open',
         LAI_4ring = 'LAI 4Ring',
         LAI_5ring = 'LAI 5Ring',
         Trans_Dir = 'Trans Dir',
         Trans_Dif = 'Trans Dif',
         Trans_Tot = 'Trans Tot',
         SF_Dir = '% Trans Dir',
         SF_Dif = '% Trans Dif',
         GSF = '% Trans Tot') %>%
  select(photo, Canopy_open_perc, Site_open_perc, LAI_4ring, LAI_5ring, Trans_Dir, Trans_Dif, Trans_Tot, SF_Dir, SF_Dif, GSF) %>% 
  filter(photo != "00AARegistration.bmp" & photo != "00_Registration.bmp") %>%
  separate(photo, into = c("sign", "val", "Pond", "Site", "T", "Ext")) %>%
  filter(sign == "n" | sign == "p") %>%
  mutate(sign = ifelse(sign == "n", -1, 1), val = as.numeric(val)) %>%
  mutate(Exp = sign * val) %>%
  select(-sign, -val, -"T", -Ext) %>%
  mutate(Camera = "DSLR",
         Resolution = "Full",
         Contrast = 0,
         Method = "Hemi")
  

YP_GLA_DSLR_fisheye <- read_delim("./data/raw/YP2020_data/YP2020_DSLRfisheye_GLA.sum", ";", escape_double = FALSE, trim_ws = TRUE) %>%
  rename(photo = 'User Field 1', 
         Canopy_open_perc = '% Cnpy Open',
         Site_open_perc = '% Site Open',
         LAI_4ring = 'LAI 4Ring',
         LAI_5ring = 'LAI 5Ring',
         Trans_Dir = 'Trans Dir',
         Trans_Dif = 'Trans Dif',
         Trans_Tot = 'Trans Tot',
         SF_Dir = '% Trans Dir',
         SF_Dif = '% Trans Dif',
         GSF = '% Trans Tot') %>%
  select(photo, Canopy_open_perc, Site_open_perc, LAI_4ring, LAI_5ring, Trans_Dir, Trans_Dif, Trans_Tot, SF_Dir, SF_Dif, GSF) %>% 
  filter(photo != "00AARegistration.bmp" & photo != "00_Registration.bmp" & photo != "00AARegistration3.bmp") %>%
  separate(photo, into = c("sign", "val", "Pond", "Site", "method", "T", "Ext")) %>%
  filter(sign == "n" | sign == "p") %>%
  mutate(sign = ifelse(sign == "n", -1, 1), val = as.numeric(val)) %>%
  mutate(Exp = sign * val) %>%
  select(-sign, -val, -"T", -Ext, -method) %>%
  mutate(Camera = "DSLR",
         Resolution = "Full",
         Contrast = 0,
         Method = "Fisheye")

## . . . Pixel spheres ====

YP_GLA_Pixel_full <- read_delim("./data/raw/YP2020_data/Pixel_full_GLA.sum", ";", escape_double = FALSE, trim_ws = TRUE) %>%
  rename(photo = 'User Field 1', 
         Canopy_open_perc = '% Cnpy Open',
         Site_open_perc = '% Site Open',
         LAI_4ring = 'LAI 4Ring',
         LAI_5ring = 'LAI 5Ring',
         Trans_Dir = 'Trans Dir',
         Trans_Dif = 'Trans Dif',
         Trans_Tot = 'Trans Tot',
         SF_Dir = '% Trans Dir',
         SF_Dif = '% Trans Dif',
         GSF = '% Trans Tot') %>%
  select(photo, Canopy_open_perc, Site_open_perc, LAI_4ring, LAI_5ring, Trans_Dir, Trans_Dif, Trans_Tot, SF_Dir, SF_Dif, GSF) %>% 
  filter(photo != "00AARegistration.bmp" & photo != "00_Registration.bmp" & photo != "00AARegistration3.bmp") %>%
  separate(photo, into = c("Pond", "Site", "T", "Ext")) %>%
  select(-"T", -Ext) %>%
  mutate(Camera = "Pixel",
         Resolution = "Full",
         Exp = 0,
         Method = "Hemi",
         Contrast = 0)

YP_GLA_Pixel_full_contrast <- read_delim("./data/raw/YP2020_data/Pixel_full_contrast05_GLA.sum", ";", escape_double = FALSE, trim_ws = TRUE) %>%
  rename(photo = 'User Field 1', 
         Canopy_open_perc = '% Cnpy Open',
         Site_open_perc = '% Site Open',
         LAI_4ring = 'LAI 4Ring',
         LAI_5ring = 'LAI 5Ring',
         Trans_Dir = 'Trans Dir',
         Trans_Dif = 'Trans Dif',
         Trans_Tot = 'Trans Tot',
         SF_Dir = '% Trans Dir',
         SF_Dif = '% Trans Dif',
         GSF = '% Trans Tot') %>%
  select(photo, Canopy_open_perc, Site_open_perc, LAI_4ring, LAI_5ring, Trans_Dir, Trans_Dif, Trans_Tot, SF_Dir, SF_Dif, GSF) %>% 
  filter(photo != "00AARegistration.bmp" & photo != "00_Registration.bmp" & photo != "00AARegistration3.bmp") %>%
  separate(photo, into = c("Pond", "Site", "c", "T", "Ext")) %>%
  select(-"T", -"c", -Ext) %>%
  mutate(Camera = "Pixel",
         Resolution = "Full",
         Exp = 0,
         Method = "Hemi",
         Contrast = 0.05)


YP_GLA_Pixel_lo <- read_delim("./data/raw/YP2020_data/YP2020_Pixel_lo_GLA.sum", ";", escape_double = FALSE, trim_ws = TRUE) %>%
  rename(photo = 'User Field 1', 
         Canopy_open_perc = '% Cnpy Open',
         Site_open_perc = '% Site Open',
         LAI_4ring = 'LAI 4Ring',
         LAI_5ring = 'LAI 5Ring',
         Trans_Dir = 'Trans Dir',
         Trans_Dif = 'Trans Dif',
         Trans_Tot = 'Trans Tot',
         SF_Dir = '% Trans Dir',
         SF_Dif = '% Trans Dif',
         GSF = '% Trans Tot') %>%
  select(photo, Canopy_open_perc, Site_open_perc, LAI_4ring, LAI_5ring, Trans_Dir, Trans_Dif, Trans_Tot, SF_Dir, SF_Dif, GSF) %>% 
  filter(photo != "00AARegistration.bmp" & photo != "00_Registration.bmp" & photo != "00AARegistration3.bmp") %>%
  separate(photo, into = c("Pond", "Site", "Contrast", "lo", "T", "Ext")) %>%
  mutate(Contrast = ifelse(Contrast == "c", 0.05, 0)) %>%
  select(-"T", -lo, -Ext) %>%
  mutate(Camera = "Pixel",
         Resolution = "Low",
         Exp = 0,
         Method = "Hemi")

YP_GLA_Pixel_fisheye <- read_delim("./data/raw/YP2020_data/YP2020_Pixelfisheye_GLA.sum", ";", escape_double = FALSE, trim_ws = TRUE) %>%
  rename(photo = 'User Field 1', 
         Canopy_open_perc = '% Cnpy Open',
         Site_open_perc = '% Site Open',
         LAI_4ring = 'LAI 4Ring',
         LAI_5ring = 'LAI 5Ring',
         Trans_Dir = 'Trans Dir',
         Trans_Dif = 'Trans Dif',
         Trans_Tot = 'Trans Tot',
         SF_Dir = '% Trans Dir',
         SF_Dif = '% Trans Dif',
         GSF = '% Trans Tot') %>%
  select(photo, Canopy_open_perc, Site_open_perc, LAI_4ring, LAI_5ring, Trans_Dir, Trans_Dif, Trans_Tot, SF_Dir, SF_Dif, GSF) %>%
  filter(photo != "00_Reg_6049.bmp") %>%
  separate(photo, into = c("Pond", "Site", "T", "Ext")) %>%
  select(-"T", -Ext) %>%
  mutate(Camera = "Pixel",
         Resolution = "Mid",
         Exp = 0,
         Method = "Fisheye",
         Contrast = 0.05)

## . . Combined ====

YP_GLA_all <- YP_GLA_Pixel_full %>% bind_rows(YP_GLA_Pixel_full_contrast,
                                YP_GLA_Pixel_lo,
                                YP_GLA_DSLR_main,
                                YP_GLA_DSLR_fisheye,
                                YP_GLA_Pixel_fisheye) %>%
  mutate(Site = paste0(Pond, "_", Site), Location = "YP2020") %>%
  select(-Pond)

YP_GLA_all %>% group_by(Camera, Exp, Resolution, Contrast, Method) %>% tally() # Missing one values from DSLR Exp 3.

## . YP combined ====

YalePreserve <- left_join(
  YP_GLA_all,
  YP_Hpix_all,
  by = c("Camera", "Exp", "Resolution", "Contrast", "Location", "Site", "Method")
) %>%
  left_join(
    YP_GLA_all %>% filter(Camera == "DSLR" & Exp == 0 & Method == "Hemi") %>%
      mutate(stdGSF = GSF) %>%
      select(stdGSF, Site),
    by = c("Site")
  ) %>%
  mutate(Site = fct_reorder(Site, stdGSF))

## Combined dataset ====

YP_RS <- bind_rows(Rockstock, YalePreserve)

saveRDS(YP_RS, "./data/processed/YP_RS.rds")
```
