## Supplementary figures and images for "Evaluation of hemispherical photos extracted from smartphone spherical panorama images to estimate canopy structure and forest light environment"

### RS017_DSLRHP.png

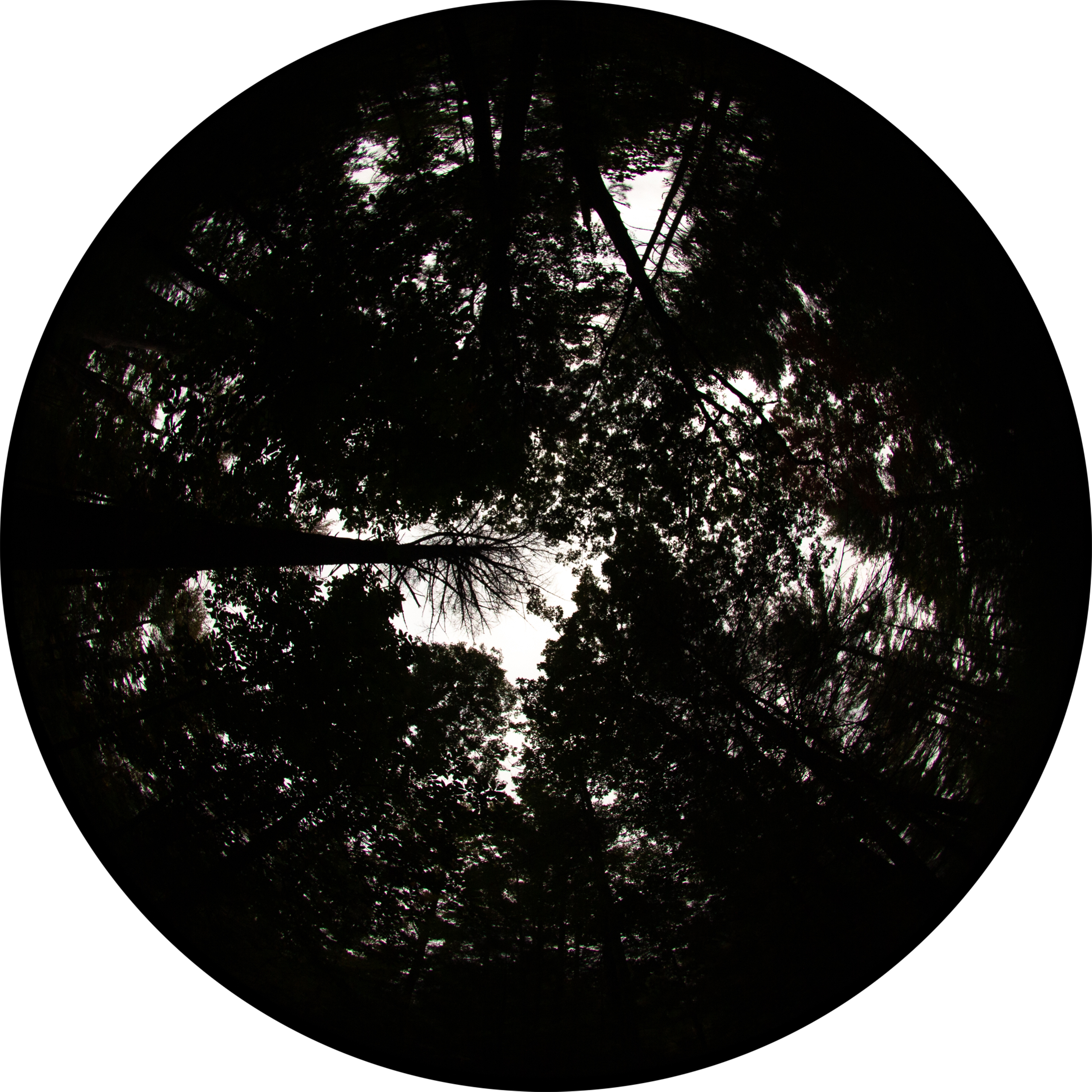

### RS017_SSPHP.png

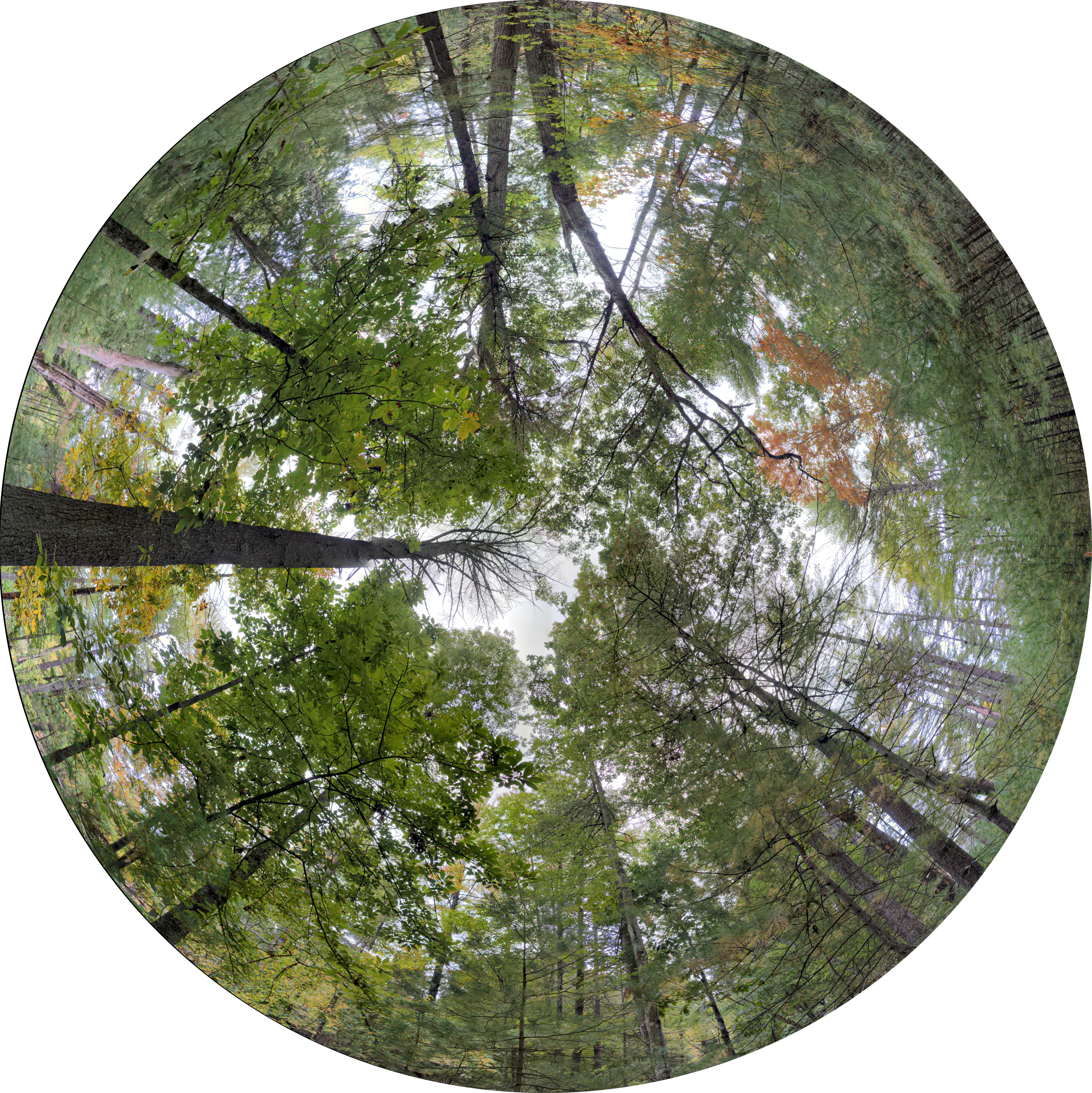
